# Transcriptome and translatome profiling and translational network analysis during seed maturation reveals conserved transcriptional and distinct translational regulatory patterns

**DOI:** 10.1101/778001

**Authors:** Bing Bai, Sjors van der Horst, Nicolas Delhomme, Alexander Vergara Robles, Leónie Bentsink, Johannes Hanson

## Abstract

Seed maturation is an important plant developmental process that follows embryo development. It is associated with a series of physiological changes such as the establishment of desiccation tolerance, seed longevity and seed dormancy. However, the translational dynamics associated with seed maturation, especially its connection with seed germination remains largely elusive. Here transcriptome and translatome profiling were performed during seed maturation. During seed maturation we observed a gradual disappearance of polysomes and a relative increase of monosomes, indicating a gradual reduction of global translation. Comparing the levels of polysomal associated mRNAs with total mRNA levels showed that thousands of genes are translationally regulated at early sates of maturation, as judged by dramatic changes in polysomal occupancy. By including previous published data from germination and seedling establishment, a translational regulatory network: SeedTransNet was constructed. Network analysis identified hundreds of gene modules with distinct functions and transcript sequence features indicating the existence of separate translational regulatory circuits possibly acting through specific regulatory elements. The regulatory potential of one such element was confirmed in vivo. The network identified several seed maturation associated genes as central nodes, and we could confirm the importance of many of these hub genes with a maturation associated seed phenotype by mutant analysis. One of the identified regulators an AWPM19 family protein PM19-Like1 (PM19L1) was shown to regulate seed dormancy and longevity. This putative RBP also affects the transitional regulation of one its, by the SeedTransNet identified, target mRNAs. Our data shows the usefulness of SeedTransNet in identifying regulatory pathways during seed phase transitions.

## Introduction

Higher plant seed development is generally divided into two phases, embryogenesis and seed maturation. Embryogenesis starts from double-fertilization occurring in the ovule and ends at heart stage when all the seed structures are defined. The cotyledon stage follows during which the embryo fills up the embryo sac and the stage ends with cell division arrest at which starts seed maturation. During this period metabolic reserves such as sugars, proteins and lipids are deposited in the endosperm and embryo (Baud et al., 2002; Baud et al., 2008) and the seed dry weight increases until reaching physiological maturation. Following this, orthodox seeds undergo desiccation drying during which metabolism gradually ceases and seed longevity is established. Eventually, the seed enters a dormant state preventing germination until the dormancy is fully released. A subset of mRNAs, produced during seed maturation, are stored in the form of messenger ribonucleoprotein (mRNP) complexes (Peumans et al., 1979), retain their functions during dry storage and are translated upon imbibition (Hughes and Galau, 1989; Comai and Harada, 1990; Bai et al., 2019). This group of stored mRNAs include mRNAs encoding ribosomal proteins, which are essential components of translation machineries (Beltran-Pena et al., 1995; Bai et al., 2017).

Regulation of translation provides an important way of determining final cellular protein levels. Average ribosome occupancies on the transcript is an optimal proxy for translation output, independent of the local variation in translation speed (Li et al., 2014). Recently, the study on the mRNAs associated with polysomes in seeds have shown distinct translational regulatory activities (Layat et al., 2014; Basbouss-Serhal et al., 2015; Bai et al., 2017). Analysis of the mRNAs that are under translational regulation have revealed several different sequence features that correlate with translational control indicating that several translational regulatory mechanisms operate during seed germination. Moreover, an imbalanced production of protein complex’ components will lead to its misfolding or degradation. Thus, protein complex components must be translated in a coordinated manner in order to efficiently allocate resources.

Gene network inference analysis allows for the functional prediction of uncharacterized genes using the guilt-by-association principle (Wolfe et al., 2005), which postulates that genes sharing similar expression profile tend to have similar functions or be involved in the same molecular pathways or functions. Common approaches rely on the clustering of gene expression profiles using a pair-wise grouping strength parameter such as a correlation coefficient followed by a hard or soft thresholding to define the network vertices. This strategy has been widely applied in different biological context and has facilitated the generation of hypothesis on gene functions and gene-gene associations (Angelovici et al., 2009; Bassel et al., 2011; Verdier et al., 2013; Costa et al., 2015; Silva et al., 2016). However, such clustering approaches disregard the edge directionality as well as the non-linearity of the regulatory cascades and are susceptible to overfitting, resulting in high network complexities; hence unsuitable for network re-construction, at least without additional assumptions (Margolin et al., 2006). Over the years, a plethora of algorithms using different approaches, *e.g.* mutual-information (Margolin et al., 2006; Faith et al., 2007b; Meyer et al., 2007; Altay and Emmert-Streib, 2010; Zhang et al., 2013), Bayesian (Friedman et al., 2000; Yu et al., 2004), graphical gaussian model (Schafer and Strimmer, 2005; Opgen-Rhein and Strimmer, 2007) or random forests based algorithm (Huynh-Thu et al., 2010) have been developed to construct gene regulatory networks (GRNs). The construction of GRNs is generally based on the calculation of the pairwise confidence scores of every edge between gene pairs (the nodes), followed by a score-based selection of edges within the full network to infer its topology, resulting in a putative causal network. More recently, Marbach *et al.* (2012), demonstrated that aggregating several gene network inferences (GNIs) in a single meta-network outperformed every individual GNI; the so-called “crowd knowledge”. This approach has recently been implemented in the *seidr* utility (Schiffthaler et al., 2018).

The seed maturation period is of vital importance for the development of viable seeds (Goldberg et al., 1994). Transcriptional events during this period are well characterized (Le et al., 2010; Yi et al., 2019). However, how translation is regulated is not known. In the current study, we investigated the translatome dynamics during seed maturation and integrated it in network analysis with our previously published translatome data from seed germination and seedling establishment (Bai et al., 2017). We discovered that genes are extensively translationally regulated during early seed maturation while late maturation is primarily regulated on the transcriptional level. Our combined network analysis identified several regulatory modules with genes sharing both function and regulatory elements. We also identified a new putative translational regulator, PM19L1, and confirmed its regulatory potential *in vivo* by mutant analysis.

## Results

### Ribosome profiles during seed maturation

To investigate translational regulation during seed maturation, ribosomal profiling was performed on maturing Arabidopsis seeds at specific developmental stages; 9-12 days after flowering (12 DAF), 13-15 DAF (15 DAF), 16-18 DAF (18 DAF) and 19-20 DAF (20 DAF). At 12 DAF, ribosomes were mainly present in the polysome form and only small peaks in monosome region were visible (Figure 1A). At 15 DAF, the total amount of ribosomes, especially that of large polysomes increased, indicating a global increase of translation (Figure 1A, B). At 18 DAF, the monosome peak dominated, while the absorbance in the polysome region of the sucrose gradient was close to background. No major changes could be noted between 18 and 20 DAF. During the maturation the overall ribosome abundance increased to until 15 DAF and thereafter decreased (Figure 1B).

**Figure 1.**
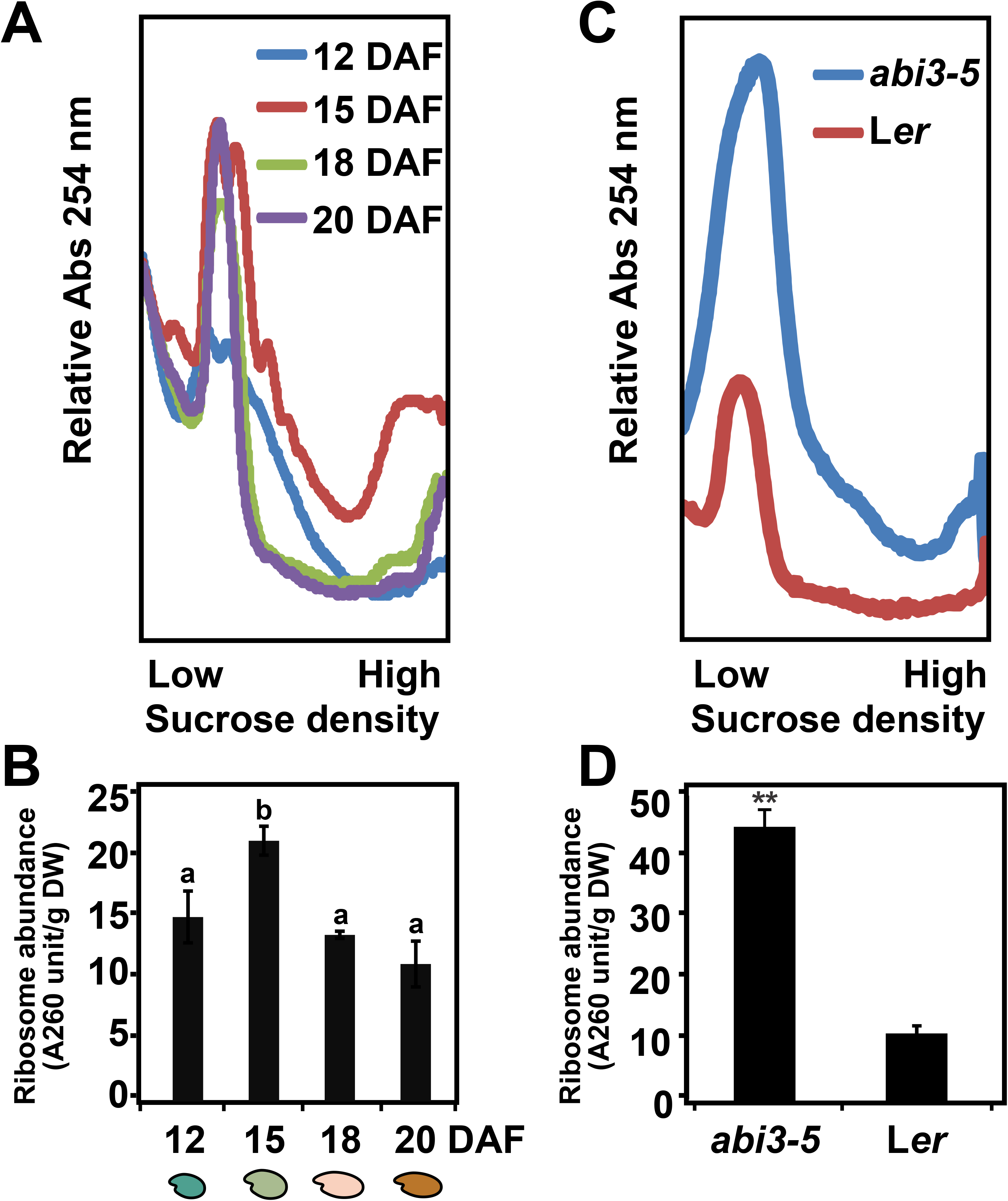
Ribosome dynamics during seed maturation. **(A)** Ribosome profile from four developmental stages during seed maturation, 12, 15, 18 and 20 DAF (Days After Flowering). **(B)** Ribosome abundance at four developmental stages during seed maturation. **(C)** Comparison of ribosome profile of the *abi3-5* mutant and wt seeds. Error bars indicate ± SD and the letters above each bar indicate the significance (*P*-value<0.05). Both seeds are harvest at the same time when wt are fully matured. The profile is based on the ribosome loading from the same seed dry weight. **(D)** Total ribosome content of wt and *abi3-5* seeds as determined by the total area under the curve in the polysome and monosome regions of the density gradient in (C). Error bars indicate ± SD and the stars indicate the significance (*P*-value<0.01).

To determine whether the accumulation of monosomes during seed maturation is developmentally regulated we analyzed the ribosomal profile of the seed maturation mutant *abscisic acid insensitive 3-5* (*abi3-5*), which displays the green seed phenotype. In *abi3-5* mutant, the embryo fully develops whereas the final maturation is impaired (Ooms et al., 1993; Sugliani et al., 2009). In the *abi3-5* seeds, a significantly higher ribosome abundance was detected compared with that of wild type Landsberg *erecta* (L*er*) (Figure 1C, D) indicating a lack of ribosome degradation, thus linking maturation to decrease in ribosomal abundance (Figure 1A).

### Transcriptional and translational dynamics during seed maturation

To investigate the translational regulation during seed maturation, total (T) and polysomal (P) mRNA was isolated for the four maturation stages and hybridized to microarrays. In total, 20,428 genes were expressed in any of the developmental stages (Supplemental Table 1). The number of genes associated with changed transcript levels decreased during seed development with thousands of genes differentially expressed at early seed maturation (12-15 DAF) and only a few hundred from 15-18 DAF. At the end of seed maturation (from 18-20 DAF) no significant transcriptional changes were identified. 29% (3,229 genes) of the genes shown to be transcriptionally regulated were also identified as differentially expressed during maturation in another study (Angelovici et al., 2009), partly validating our results (Fig. S1A). The transcriptional changes during maturation are conserved between Arabidopsis and Medicago since largely overlapping transcriptional changes were detected in data from *Medicago truncatula* seed maturation (Verdier et al., 2013) (Fig. S1B-G).

To identify genes translationally regulated during seed maturation, the change in polysome occupancy (PO) defined by the ratio between the polysomal mRNA and total mRNA intensities of each stage was determined. Large PO changes were identified between 12-15 DAF with significant more changes in the positive direction, Figure 2A (2144 genes increased in PO in contrast to 1087 genes with decreased PO). Analog with other translational reprogramming events during seed germination (Bai et al., 2017), we term this change Maturation Translational Shift (MTS). At the later stages during maturation no significant changes in PO was identified (Supplemental Table 2). This finding is supported by the Principal Component Analysis (PCA) where the relation between the polysome samples and the total RNA samples are different in 12 DAF samples compared to the relation at the other points (Figure 2B). In general, the translationally up-regulated genes were expressed to relative low levels and similarly weakly associated to the polysomes in 12 DAF. At 15 DAF these genes were associated to polysomes at higher than average levels and this trend persisted until the later stages, and thus a PO increase preceded the relative transcriptional change (Figure 2C). However, as only relative expression levels can be determined by microarrays it is possible that the increased relative level is caused by decreased mRNA stability or general decrease of transcription. The opposite pattern was shown for the down-regulated genes. These genes were generally associated with polysomes at 12 DAF and detached from polysome at 15 DAF. Several of the translational regulated genes identified here were under translational control during seed germination and in response to hypoxia stress as well (Supplemental Figure 2A, B). Less overlap was identified with translational regulation as a result of sucrose starvation or feeding (Supplemental Figure 2C, D). GO analysis for the translationally regulated genes revealed that diverse molecular functions and biological processes were under translational control during seed maturation (Supplemental Table 3). The translationally up regulated genes were enriched for GO terms including flavonoid biosynthesis, vegetative to reproductive phase transition, chromatin assembly, ABA metabolic process and signaling. Translationally down regulated genes were enriched for functions including glycolytic process, fatty acid biosynthesis, gluconeogenesis, photosynthesis and stress response.

**Figure 2.**
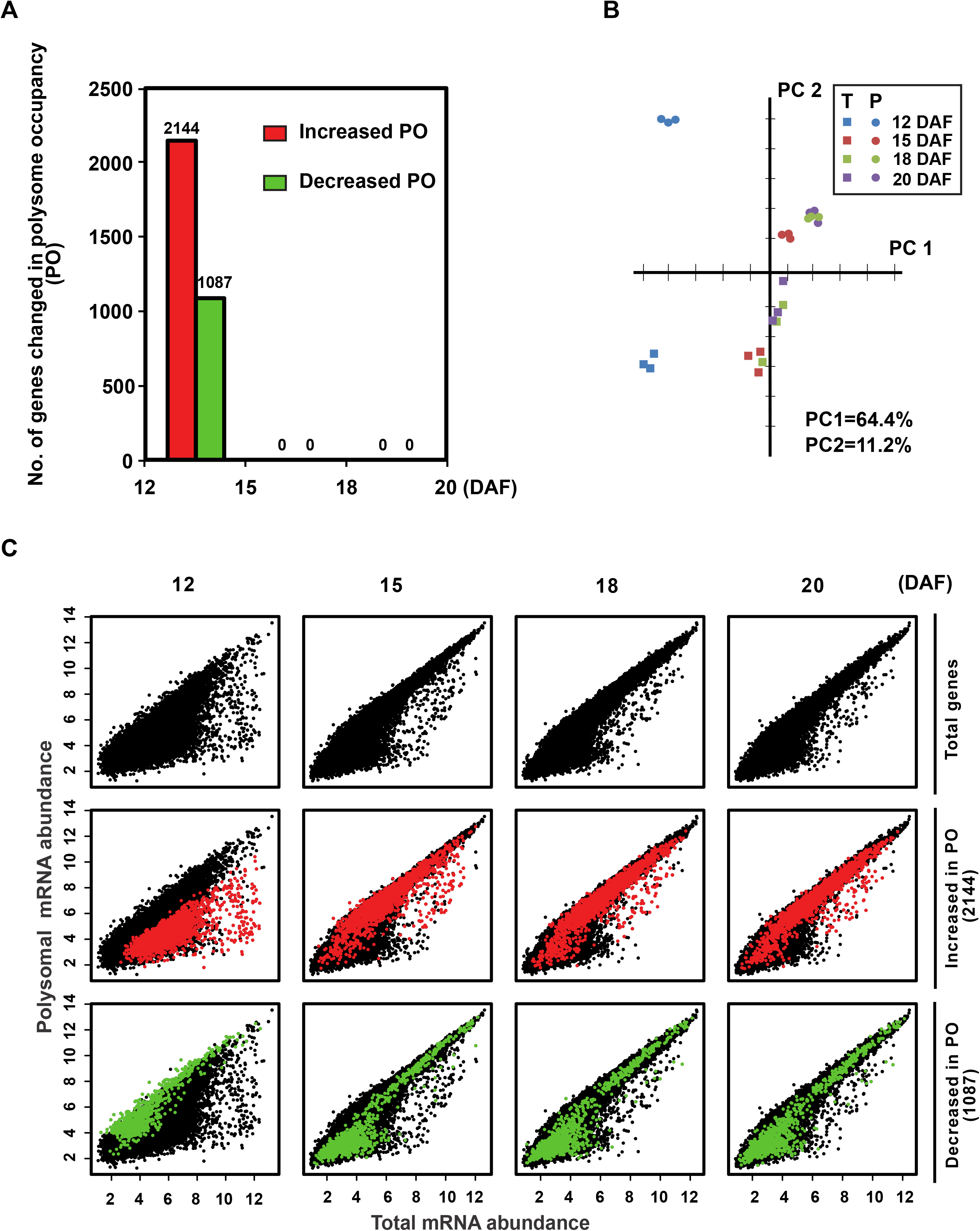
Translational Shift during Arabidopsis seed maturation. A. The number of genes with changed polysome occupancy (PO) during seed maturation. Seed were harvested 12, 15, 18 and 20 days after flowering (DAF). The increase or decrease in PO are indicated in red and green bars respectively with gene number above the bars. B. Principal component (PC) analysis of the total RNA (T) and polysomal RNA (P). normalized expression levels during seed maturation. Individual biological replicates are highlighted in the same color. The sample of total RNA and polysomal RNA are indicated as circle and square respectively. The different colors represent different time point during seed maturation. The first two components PC1 and PC2 explain 64.4% and 11.2% of the total variation, respectively. C. Relation between normalized abundance of genes in the polysomal and total RNA preparations respectively. The genes associated with significant changed PO between 12-15 DAF (MTS genes) are indicated in red (increased PO) or green (decreased PO) respectively.

To further classify the translationally regulated genes, we clustered these genes based on their transcriptional and polysomal changes from 12-15 DAF (Supplemental Figure 3). In total 12 translational groups were identified which exhibited diverse modes of translational regulation and distinct molecular functions (Supplemental Table 3). The different patterns suggest that several distinct modes of translational regulation during seed maturation.

### Transcript features correlate with translational regulation during seed maturation

In order to identify shared sequence features for the translational regulated genes that may influence the translation such as UTR length and GC content (Kawaguchi and Bailey-Serres, 2005), sequence analysis was performed on the transcripts that changed in PO during seed maturation. Transcripts that increased in PO had generally longer CDS and the opposite was observed in the transcripts with reduced PO (Supplemental Figure 4A and C). A significant higher GC content was detected in the CDS region of transcripts with increased PO (Supplemental Figure 4B and D). No significant difference was found for GC3 (Guanine and Cytosine content in the third codon position; synonymous sites) and effective number of codons (Nc) (Supplemental Figure 4E), indicating codon bias is not separating the translationally up or downregulated regulated mRNAs during seed maturation. To test if differently regulated mRNAs have different overall mRNA secondary structure was tested by comparing an experimentally derived structural score defined by Li *et al*. for the different set of mRNAs (Li et al., 2012). The score is an indicator of potential transcript complexity in which high structure scores are equivalent to more double stranded (ds) structures at a certain position in a transcript compared to regions with lower score. In general, the translationally up-regulated genes had lower than average structure score (Supplemental Figure 4F), especially in the coding region. Translational down-regulated genes were in this analysis showing higher than average scores especially at the end of the 5’leaders and in the coding region. The transcripts with highest structure scores, were enriched for photosynthesis and thylakoid GO categories (Supplemental Table 4). Probably the translational regulation of these photosynthesis related genes during seed maturation is mediated the mechanism involving the mRNA secondary structure.

### Translational dynamics display functional association during seed maturation and germination

Previously, two distinct translational regulation shifts (termed Hydration Translational Shift, HTS, and Germination Translational Shift, GTS, respectively) have been identified during seed germination (Bai et al., 2017). In this study we have identified one single stage, at the early stage of seed maturation, where translation is regulated; the Maturation Translational Shift, MTS. Plant development can be seen as one trajectory starting with the embryogenesis until the mature plant, this development is interrupted by a quiescent period in the seed. Here, we treated the combined seed data, from early maturation to seedling development, as a single time course. Genome-wide polysome occupancy (PO) profiles of the combined dataset were generated by pheatmap tool (Kolde, 2018) as proxy for gene translation efficiency across seed maturation and germination. The obtained PO profile generated displayed a dynamic pattern during seed maturation and germination (Figure 3). The PO dynamics was represented by clusters in which the transcripts showed similar translational dynamics (Figure 3, Supplemental Table 5). Cluster 1, 3, 7, 12 represented transcripts with enhanced in PO during seed maturation and reduced PO during seed germination; Cluster 5, 6, 10, 11 contained transcripts with reduced PO during seed maturation and different in dynamics during seed germination; Cluster 2, 5 contained by transcripts with changing PO only during seed maturation. Cluster 8 and 13 represent genes with consistently changing PO during seed maturation as well as during seed maturation and germination. As expected, transcripts from *e.g*. cluster 3 are enriched for transcripts classified for the biological process “water deprivation” and “embryo development ending in seed dormancy” enhanced in PO during maturation which are important process during seed maturation while cluster 4 and 11 enriched in transcripts involved in post-embryonic growth and seed germination, which are increased in PO at different stage during seed germination (Supplemental Table 6). The correlation between distinct regulatory patterns and discrete gene function strengthen the biological relevance translational regulation during the studied developmental transition.

**Figure 3.**
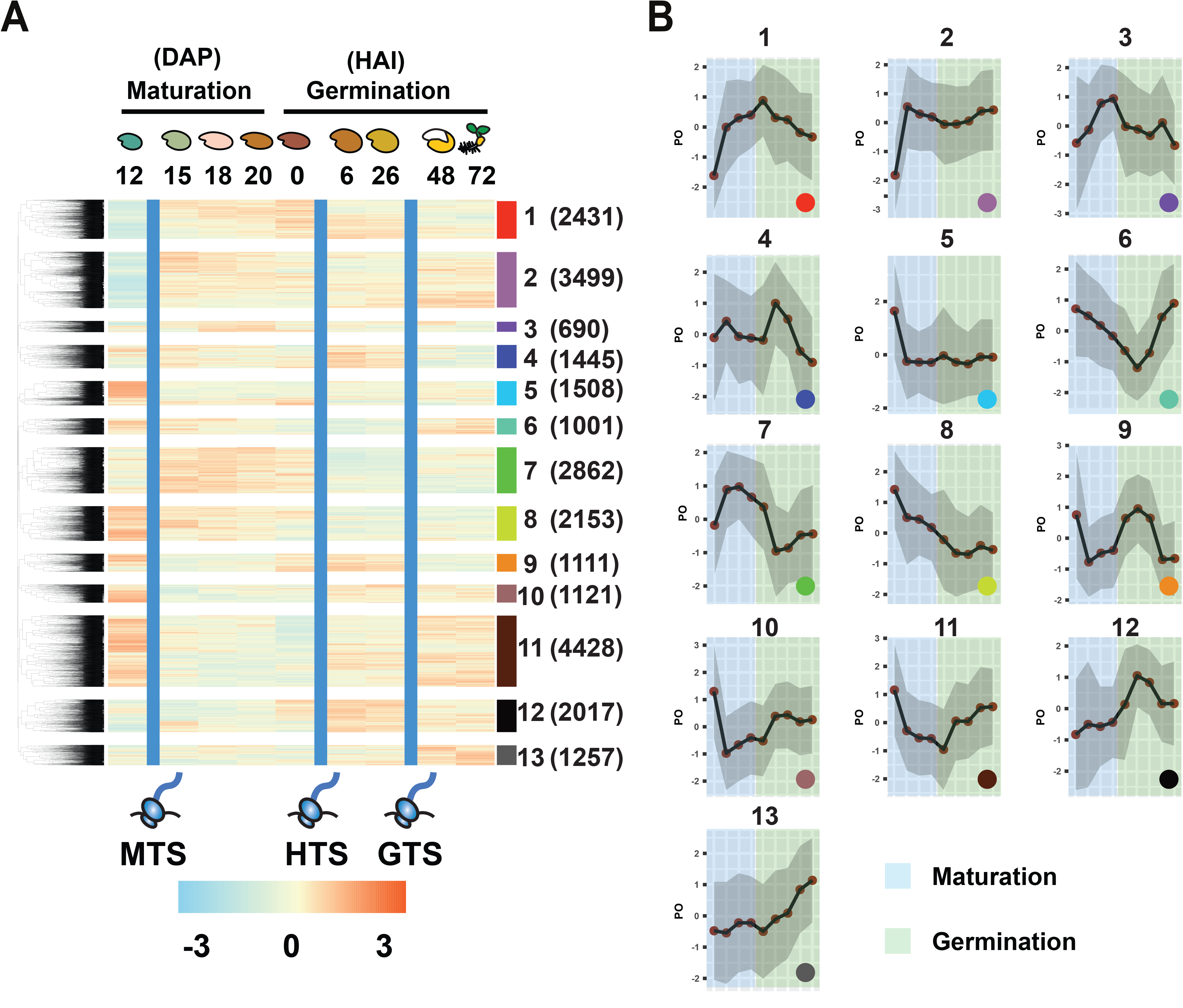
Dynamics of polysome occupancy during seed maturation and germination. A. Heatmap and hierarchical clustering of polysome occupancy during seed maturation and germination. The time course including 12, 15, 18, 20 days after flowering (DAF) from this study and after-ripened dry seeds (0) as well as seeds imbibed for 6, 26, 48 72 hours (HAI) from the study from (Bai et al., 2017). The average PO intensity of three biologicals replicates are plotted. Heat red yellow and blue represent the high, media and low PO intensity respectively. The number of the clusters is represented with different colors with the number of genes in each cluster in the bracket. The blue bars that separate the developmental stages represent three translational shifts (Maturation Translational Shift (MTS), Hydration Translational Shift (HTS) and Germination Translational Shift (GTS) respectively). B. Trend plot of PO levels across the whole time-course for each of the corresponding clusters identified in A. The trend plots are separated for the developmental stages by seed maturation and germination. The color of each cluster corresponds to the clusters identified from A.

### Translational network reveals translational association and distinct community features with specific functions during seed maturation and germination

The cluster analysis describes the translational pattern but neither interaction nor directionality, thus we cannot predict regulators nor their targets from this analysis. As our data set represents a time-course it can be used to determine regulatory pathways. We therefore employed a directed regulatory network approach using the Seidr tool Schiffthaler et al., (2018). The output of Seidr is a weighted gene-gene association matrix based on the translational profile including the resource and target of every edge, their interaction strength, directionality (which is either directional or undirectional) and ranks for each of the inference algorithms. Based on the network topology (measured by its scale free transitivity and average clustering coefficient), a stringent threshold score was devised to the select the most relevant network edges (Supplemental Figure 5, see material and method). This result in a putative gene regulatory network, which we call seed translation network (SeedTransNet), with 7873 nodes and 8740 edges, 2298 of which are directed (Figure 4A, Supplemental Table 7).

**Figure 4.**
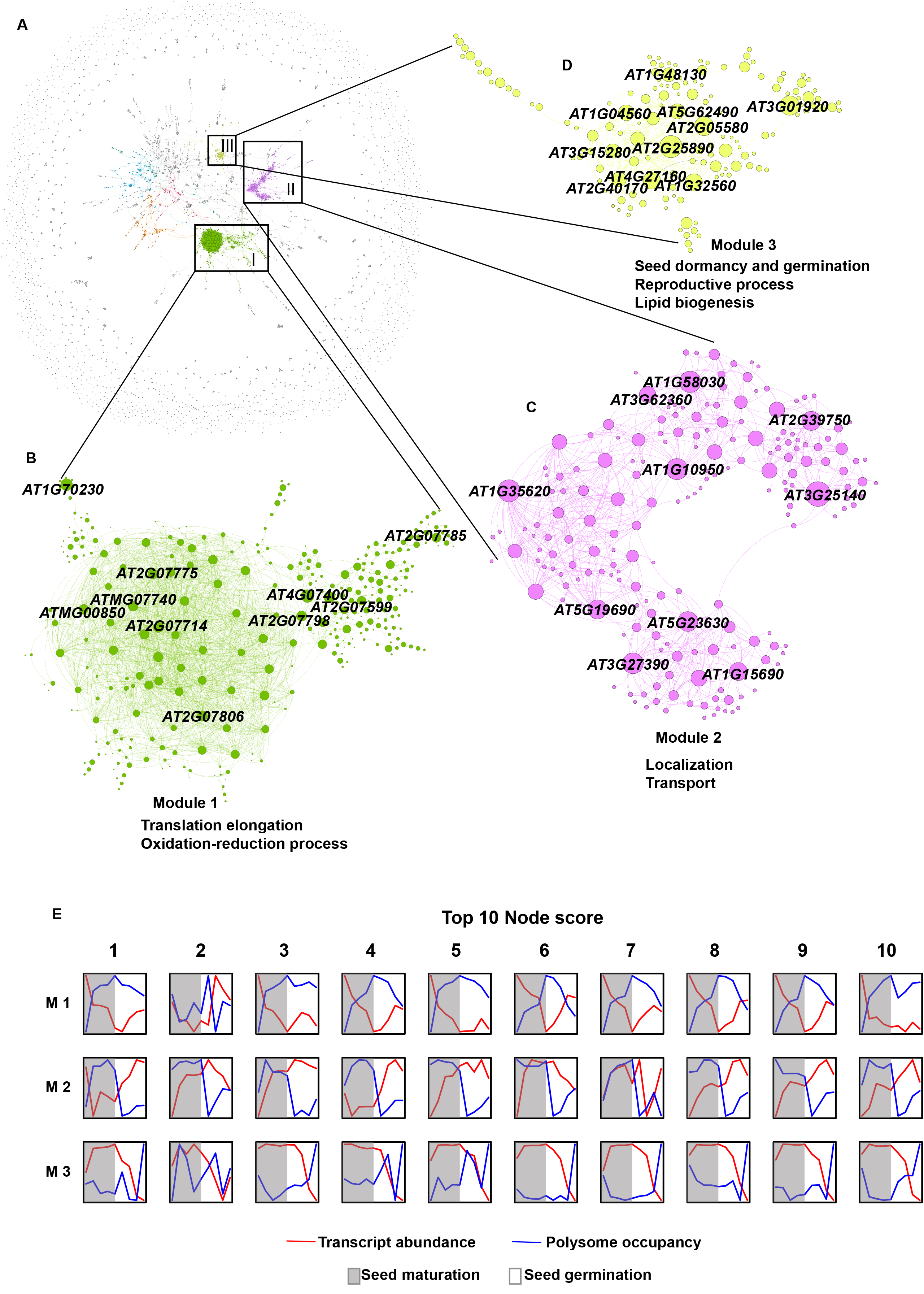
SeedTransNet, a directional gene regulatory inference network based on the combined dataset of PO during seed maturation and germination (this study and Bai et al. 2017). A. SeedTransNet with nodes representing each gene and edge representing the pairwise PO linkage. Color indicating the modules detected from the SeedTransNet. The major modules are color coded. B-D. Module topology of the Top 3 modules and their associated major GO terms. Nodes with high connectivity within each module are depicted as bigger circles. Module 1 includes 3134 genes predicting 1799 gene linkage inferences (GLI) with average module node degree of 10.78. Module 2 contains 364 genes predicting 684 GLI with average module node degree of 6.91. Module 3 is consisted of 140 genes representing 840 GLI with average module node degree of 4. E. Transcription and polysome occupancy plotted across seed maturation (grey) and germination (white). The top ten nodes from each of the three top modules are displayed.

Co-expression networks that include highly connected gene nodes could reveal functionally coherent modules and their regulators (Segal et al., 2003). We thus identified from SeedTransNet gene module (highly connected gene clusters) structure using Infomap (Rosvall and Bergstrom, 2008). This algorithm uses the probability flow of random walks on the network as proxy for devising the information flows in the “real” system informing on the network structure and node relationship. The modules identified in the SeedTransNet was organized in a hierarchical modular structure based on the Infomap results (level M1 to M5, Supplemental Table 8). Each module level represented a different level of granularity which may indicate functional hierarchy in the network. SeedTransNet is characterized by a highly modulated center core surrounded by a fragmented network ring with relatively low network density and modularity. In total 1857 modules were identified on the first level of granularity (M1) were identified, out of which 572 could be assigned with a specific biological function by GO enrichment analysis (Supplemental Table 8 and 9). The high functional modules detected indicated that SeedTransNet could be used to discover new gene translational regulatory pathways. The top three modules (Module 1-3) in the M1 granularity level which displayed visible modularity features and translational dynamics in the network were further explored for gene linkage inference (GLI) based on their high gene linkage prediction (10.46% of the total putative gene translational linkage) (Figure 4B-D, Supplemental Table 8).

Module 1 module represents transcripts, which transcription was strongly reduced during seed maturation but activated during seed germination while the opposite dynamics was observed for their polysome occupancy (PO) (Figure 4E). Module 1 is enriched for mitochondrial genes (70/334), especially involved in translation and mitochondria electron transport (Supplemental Table 8), indicating that mitochondria genes are likely under stringent translational control during seed maturation and germination.

Module 2 contains transcripts that showed high level of PO during seed maturation are similarly to the Module 1 while polysome dissociated at early seed imbibition and recruited during late stage of seed germination, different from Module 1 (Figure. 4E). Gene enrichment analysis identified diverse gene functions involved in transport and protein glycosylation (Figure 4C and E, Supplemental Table 9). Among the abundant transporters identified was the well-known germination regulator-vacuolar Ca^2+^ activated channel protein gene TPC1, which governs abscisic acid (ABA: phytohormone that inhibit seed germination)-sensitivity during seed germination and thus the *tpc1* mutant is not response to ABA during seed germination (Peiter et al., 2005). Protein glycosylation is important for cell wall synthesis and maturation, which is also an active process during seed germination due high levels of cell division and elongation.

Module 3 displayed a low PO during seed maturation while the transcript accumulated. These accumulated transcripts showed sharply reduction in abundance upon seed germination when the PO drastically increased (Figure. 5E). These transcripts are possibly seed stored mRNAs that accumulate in seeds and are activated for translation at different stages during seed germination. (Figure 4E). GO enrichment analysis show that genes in this module were mainly associated with seed development, post-embryonic development, seed dormancy and germination and lipid biogenesis including large amount of Late Embryo Abundant (Lea) protein genes, Oleosin genes and seed storage protein genes (Figure 4D, Supplemental Table 9). LEA proteins are distinctly identified in Module 3 (Supplemental Figure 6A, Supplemental Table 10). These LEA proteins were translationally linked with oleosin family protein, crucifer family seed storage protein and other proteins related to ABA, cell wall synthesis and ROS balance (Supplemental Figure 6A). The close relation of Module 3 with seed development was seen by the enriched *ABSCISIC ACID INSENSITIVE3* (*ABI3*) regulated genes (*ABI3* regulon) which one of the key regulators of seed development (Monke et al., 2012). In total 43/98 (43.9%) of the ABI3 core regulon were specifically identified in this module (Supplemental Figure 6B, C, Supplemental Table 11), indicating an association of Module 3 with down-stream targets of ABI3. This linkage was connected with genes encoding oleosin, LEA, and storage protein in SeedTransNet (Supplemental Figure 5C), indicating lipid synthesis protein, seed storage protein and late abundance protein share common feature modulated by ABI3. Above all, the network analysis identified putative hierarchical modules with distinct functions and could be used for further gene regulatory inference.

**Figure 5.**
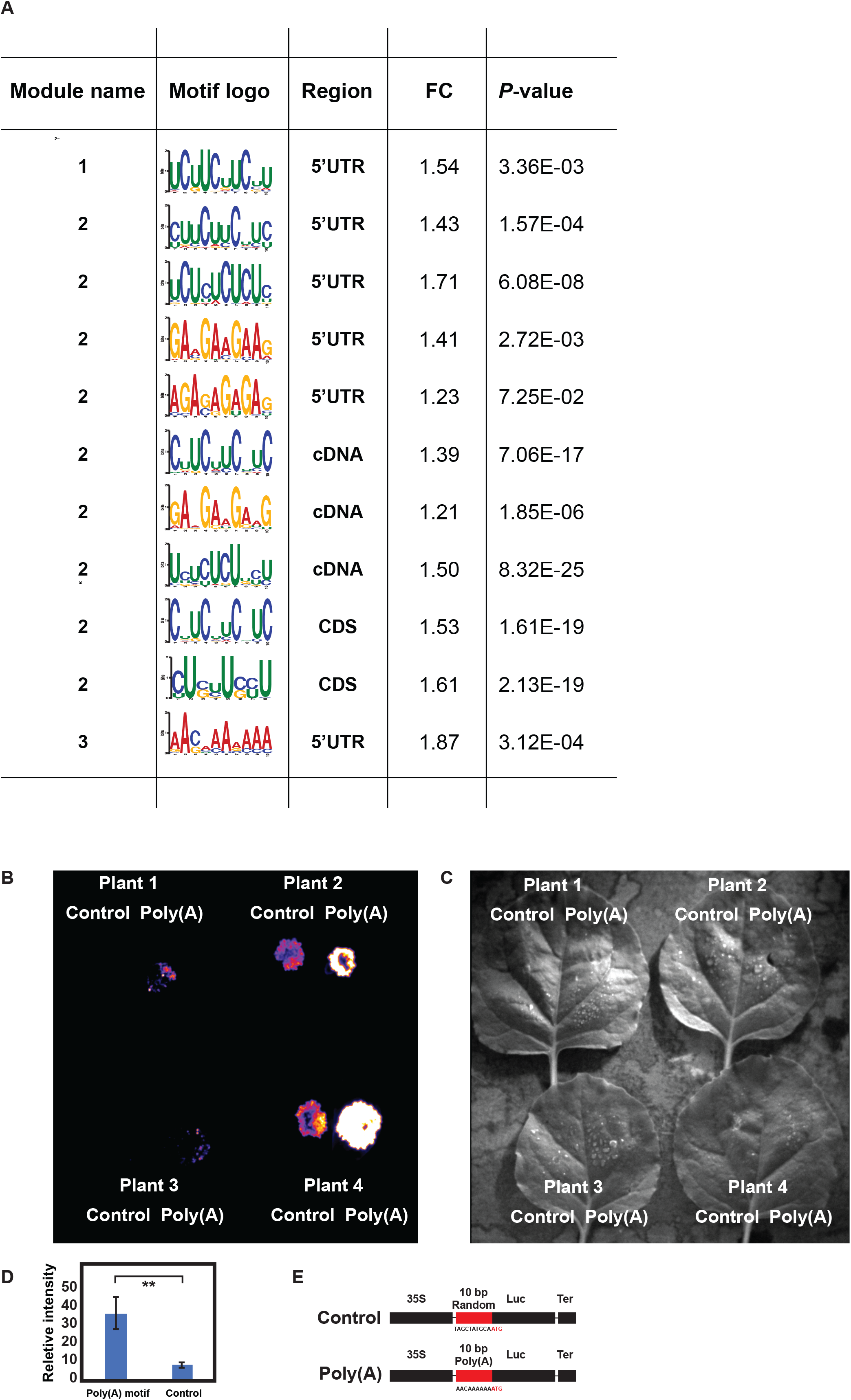
Functional analysis of cis-elements identified from SeedTransNet. A. The significantly enriched motifs detected in the Module 1-3 of the SeedTransNet. The sequence logo, motif localization, fold change (FC) enrichment compared with the background sequence (total microarray background sequence at each sequence region) and *P*-value (Fisher test) are showed. B. Luciferase signal detected 3 days after infiltration of *Nicotiana benthamiana* leaves with construct containing 35S promoter fused to luciferase gene with either the poly(A) motif (right) or a random 10 bp control sequence (left) respectively at the 5’UTR. C. Gray light image corresponding to the image in C. The position of luciferase signal on the leave discs from B is corresponding to the position in C D. The quantification of relative Luciferase activity detected 3 days after infiltration of *Nicotiana benthamiana* leaves with construct containing 35S promoter fused to luciferase gene with either the poly(A) motif (right) or a random 10 bp control sequence (left) respectively at the 5’UTR. Error bars indicate ME±SE (n= 8), stars indicate *P*-value < 0.01 (t-test). E. Schematic drawings of the control and poly(A) motifs construct used for transfection, the black bars represent 35S promoter, luciferase coding sequence and terminator. The red bar indicated the motif sequence followed by the start codon ATG for the luciferase gene.

### SeedTransNet identifies RNA motifs regulating mRNA translation

Translational regulation is often mediated through cis-elements in the target mRNAs. By comparing each mRNA sequences of each module with mRNA sequences of the whole genome, motifs specifically enriched in different regions for each module were identified (Figure 6A). Pyrimidine (UC) enriched motif were identified in 5’UTR of Module 1, while pyrimidine with different repeats (UCM and CUU) and complementary purine repeats (GA and GAA) enriched motifs were identified in the 5’UTR and cDNA sequence of Module 2. In the Module 3, a A-rich motif AACAAAAAA was enriched in the 5’UTR (Figure 6A). Interestingly, the same motif was identified in the transcripts with increased PO during early seed germination (Bai et al., 2017). A similar poly(A) motif was recently identified to play a role on gene specific translational regulation in response to pathogen by interacting with poly(A) binding proteins in Arabidopsis (Xu et al., 2017). To confirm the function of the identified poly(A) motif in modulating translation, the motif was fused to the luciferase reporter gene under the control of the 35S promoter and the construct was infiltrated into *Nicotiana benthamiana* leaves. Compared to a similar construct with a control motif of 10 bp, the poly(A) motif generated on average 3-5-fold higher luciferase signals (Figure 6C-E, Supplemental Figure 9A, B), while the abundance of *LUC*-transcript was not significantly changed (Supplemental Figure 9C). This experimentally confirms the potential role for the identified poly(A) motif in translational regulation.

**Figure 6.**
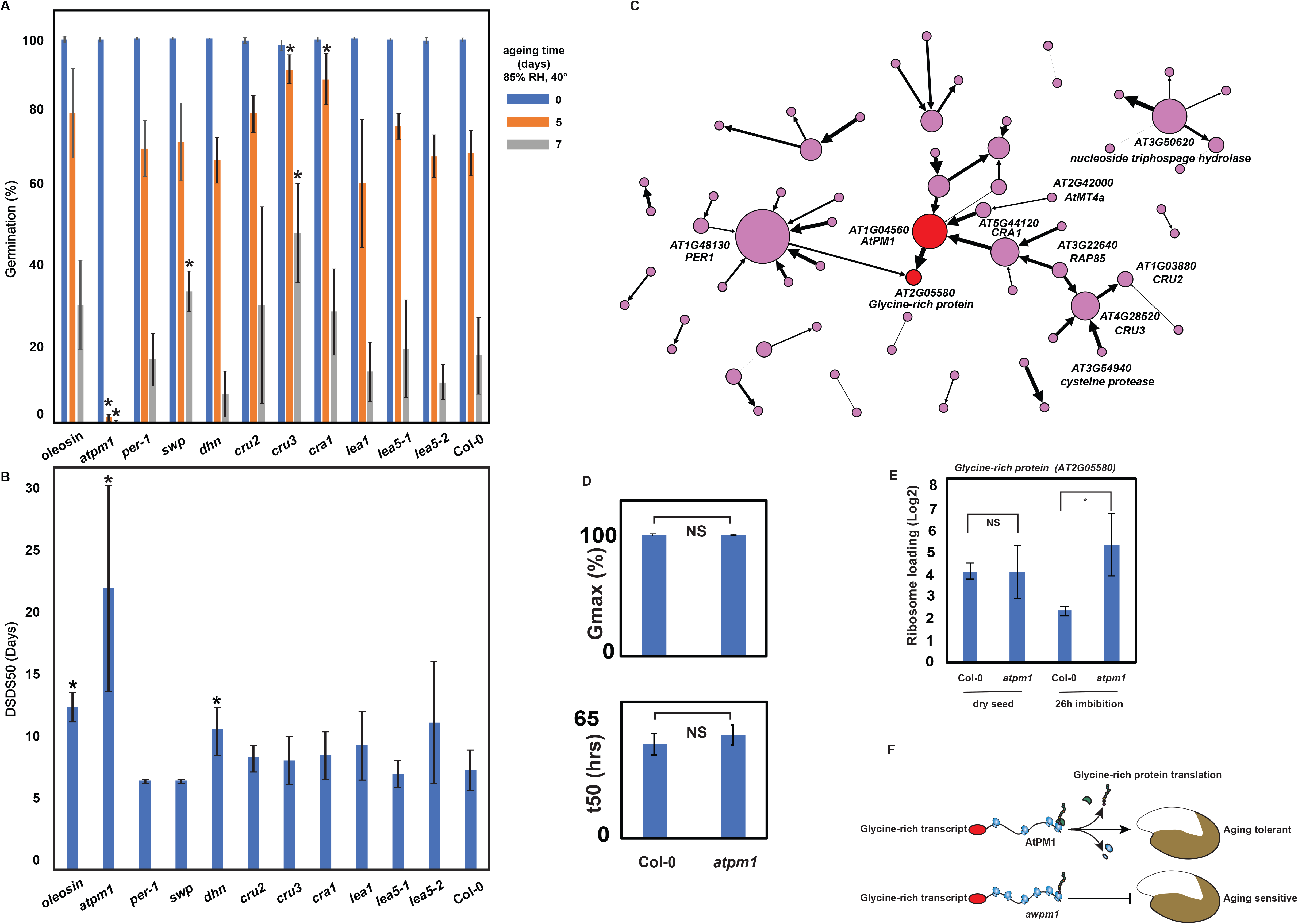
SeedTransNet regulatory network hub genes are affecting seed characteristics. A. Seed longevity phenotype of mutants of hub genes selected from M3 module. Five and seven days aging test were used for evaluating seed longevity in comparison with zero days aging. Error bars indicate ME±SE (n= 4), stars indicate *P*-value < 0.05 (t-test). B. Seed dormancy phenotype of mutants of hub genes selected from M3 module represented by DSDS50 (the days of seed dry storage required to reach 50% germination). Error bars indicate ME±SE (n= 4), stars indicate *P*-value < 0.05 (t-test). C. Close up of directional network in the M3 module centered around AT1G04560 and the target gene confirmed experimentally, AT2G05580 (both red). Strengths of the respective predictions are indicated with edge thickness. D. The maximum germination (Gmax) and time need to reach 50% germination (t50) are indicated to both the *atpm1* and Col-0 seed before the ribosome isolation. Error bars indicate ME±SE (n= 4), NS indicate no significant difference, *P*-value < 0.05 (t-test). E. Ribosome occupancy (RO) of *Glycine-Rich Protein* gene transcript change in response to imbibition in wild-type and mutant seeds. F. The hypothesized function of AtPM1 in translational regulation of its down-stream target *Glycine-Rich Protein* gene.

### Network inference hints on translational regulation during seed maturation and germination

The dependency-based property of SeedTransNet allows the identification of directional gene inference. In total, 65 directed gene inferences were predicted from the modules within the top 3 modularity ranking, among which 43 are predicted from module 3 (Supplemental Table 11). To validate the predicated inference, an arbitrary node degree cutoff of 5 was used to select the nodes that may constitute the main translational regulators. Five interactions passed this selection criteria with the five nodes were associated with embryo maturation including 1-cysteine peroxiredoxin1 (PER1), *AWPM-19-LIKE* protein (AT1G04560), *CRUICIFERINA1* (CRA1), *CRUCIFERIN2* (CRU2) and *CRUCIFERIN3* (CRU3).

We tested the influence of network hub nodes in both directional and in-directional way in Module 3 for their influence on seed maturation and germination by investigating their seed dormancy and longevity phenotypes. The mutant of one of the hub node genes encoded the AWPM-19-LIKE1 (PM19L1) protein and showed extreme sensitivity to ageing as seen by the sharp reduction of seed germination after 5 days ageing compared with 80% germination at the same ageing stage of the wildtype seeds. Similar results were found for two independent T-DNA knock-out mutants and the aging sensitivity can be complemented by over-expression of *PM19L1* driven by the 35S promoter (Figure 6A, Supplemental Figure 7). The mutant, *pm19l1,* showed significantly enhanced seed dormancy (Figure 6B) as well as shown independently by (ref. Barrero et al 2019). Taken together this indicates that the PM1L1 protein plays an essential role for seed maturation. Judging from the SeedTransNet inference, AtPM1 translationally regulated a down-stream target gene *AT2G5580*, encoding for a Glycine-Rich Protein (Figure 6C). By comparing with the public protein-protein interaction network STRING (Szklarczyk et al., 2015) co-expression network GeneMANIA (Warde-Farley et al., 2010), no direct co-expression of the two genes can be predicted while these two genes share some co-expressed gene nodes (Supplemental Figure 8A, B) such as LEA protein genes and oleosin family genes. The expression of both of *PM19L1* and *Glycine-Rich Protein* gene are highly seed specific (Supplemental Figure 8C, D). Thus, we hypothesized that PM19L1 protein may translationally regulate the *AT2G5580* gene. To validate this, ribosome associated mRNAs were isolated from the *pm19l1*in both dry seeds and 26 hours after imbibition (HAI) when seeds were completely released of dormancy and displayed seed germination dynamics as Col-0 wild-type (Figure 6D, E). In the wild-type plant PO of *AT2G5580* is decreased more than twofold upon germination. This is totally abolished in the *atpm1* mutant, experimentally indicating the importance of the PM19L1 protein for the translational regulation of AT2G5580 as predicted by SeedTransNet inference.

## Discussion

### The accumulation of monosomes during seed maturation is developmentally regulated

Seed maturation is an intricate developmental process starting when the embryo is fully developed. Seed maturation is characterized by growth arrest, storage reserve accumulation, acquisition of desiccation tolerance, and the induction of seed dormancy. In this work, we presented the first translational dynamics during seed maturation by investigating polysomal profiling at four stages during seed maturation. Seed maturation in which the seed is associated with distinct polysomal dynamics. There is high polysomes abundance during early maturation, these polysomes disassemble at the end of the seed maturation which results in mainly monosomes in the mature dry seed (Figure 1). The seeds of the severe seed maturation mutant (*abi3-5*) do not show the reduction in polysomes during seed maturation that are seen in wild type. These seeds are also unable to fully accomplish seed maturation seen by reduced longevity, dormancy and tolerance desiccation (Koornneef et al., 1982; Koornneef et al., 1984; Ooms et al., 1993). Our data strongly indicate that the disassembly of polysomes into monosomes is developmentally regulation during seed maturation.

### Translational reprogramming affects the expression pf seed maturation genes

Our data largely confirms a conserved transcriptional changes during seed maturation of different species as previously identified (Supplemental Figure 1), including these key regulators for seed longevity such as *NFXL1* and *WRKY33* (Supplemental Figure 1C) (Righetti et al., 2015) as well as dormancy related *DOG1* and *ABI3* which are known to play important role for seed dormancy, which is established during seed maturation (Supplemental Figure 1E).

The genome-wide polysome profiling indicated that translational regulation is concentrated during early phase of seed maturation (MTS) from 12-15 DAF. This translational shift is temporally correlated to seed desiccation and nuclear size reduction (Baud et al., 2002; van Zanten et al., 2011). Maturation programmed desiccation may play an important role for translational control since it is known that dehydration could influence mRNA translation (Kawaguchi et al., 2004).

Genes affected by MTS display diverse transcriptional and translational profiles (Supplemental Figure 3). This might indicate MTS genes are differently regulated. Among the translationally controlled genes are genes that have been reported as seed specific regulators eg *Abscisic Acid (ABA)–Insensitive 2* and *5* (*ABI2* and *ABI5*) (Supplemental Table 12), the mutants of which are insensitive to ABA during seed germination (Finkelstein, 1994; Rodriguez et al., 1998). The translational profiles of both *ABI2* and *ABI5* are correlated with the ABA accumulation pattern during seed maturation that peaks during early seed maturation when cell elongation is terminated (Vicente-Carbajosa and Carbonero, 2005). Besides *ABI5*, ABA synthesis and signaling factors *ABA DEFICIENT2* (*ABA2*) and *Dehydration-Responsive Element Binding Factor 2C* (*DREB2C*, AT2G40340) are translationally regulated. *STAY-GREEN2* (*SGR2*), important for regulating the chlorophyll content during seed maturation is also translationally regulated. The PO of the key seed dormancy regulator *RDO4*, the mutant of which showed reduced seed dormancy (Liu et al., 2007), is also enhanced during seed maturation. There examples implies the role of translational control of molecular events preparing seeds for the final ripening and prime the seed for germination.

### Sequence length, GC content and secondary structures correlate with changed polysomal occupancy regulation during seed maturation

Although the mechanistic understanding is not clear, occurrence of certain sequence features has been shown to correlate with translational regulation (Kawaguchi and Bailey-Serres, 2005; Vogel et al., 2010). Our sequence analysis revealed that a few sequence features are correlated with polysome association during early phase of seed maturation. Under this developmental transition, the transcripts with increased PO are longer. Genome-wide analysis has shown an inverse correlation between CDS length and translation since ribosome recycling is more efficient on shorter transcripts (Arava et al., 2003; Vogel et al., 2010; Liu et al., 2012; Fernandes et al., 2017). Our length data are similar to the CDS length feature identified from previously published data on the translational regulation upon heat stress during which transcripts with long CDS are favored for polysome association under stress condition while short transcripts are preferred at normal condition (Yanguez et al., 2013).

The translationally up regulated transcripts in our study have higher GC percentage in the CDS (Supplemental Figure 4B, D), and thus in our dataset GC-rich coding region is over-represented in transcripts up-regulated in PO indicating an adaptive role for translation during seed maturation. Same GC-enriched coding region has been shown for mRNAs in association with ribosome in rice (Zhao et al., 2017), indicating a conserved sequence feature for ribosome association.

The translationally upregulated genes during MTS has generally lower structure score in their CDS compared to downregulated or the background set (Supplemental Figure 4F). The low structure score of these transcripts may facilitate the enhanced accessibility for ribosome attachment (Li et al., 2012) or make these transcripts less prone to stalling during desiccation. Possibly, translational elongation is important factor for regulating the translational speed during seed maturation, e.g. the adaptation of the codon for the tRNA specifically expressed during development or the folding feature of the CDS, which has been also reported on mouse embryonic stem cells (Dana and Tuller, 2012).

### Network analysis reveal regulatory circuits possibly controlled by distinct sequence motives in the regulated mRNAs

By constructing a directional co-translational network, SeedTransNet, with data from consecutive developmental phases including both seed maturation and germination, we could identify functional modules with distinct functions along the whole time-course. Using this network, gene regulatory hubs and the directional regulatory cascades could be accurately predicted, as judged by later experimentally confirmation. We both confirmed regulatory hub genes to have important role during seed maturation and germination by analysis of corresponding mutants and by confirming regulator-target gene relationship using regulatory studies. Only polysomal occupancy data was used to construct SeedTransNet but the resulting module genes share both polysomal occupancy patterns and mRNA level changes (Figure 4E), indicating genes identified from the same module might be under same regulatory mechanism for their expression. In addition, we could identify regulatory elements in the mRNAs using reporter gene analysis. Therefore our network is providing a promising recourse for future crop engineering and seed quality improvement efforts. Here something about the database and the web recourse that we will provide. Interestingly, the network modules cluster genes with similar molecular function. As represented here, the top three modules showed distinctive GO functionality. Module 1 is enriched with translational factors including substantial ribosomal RNA and tRNA, corresponding to the dramatic changes in ribosome profile identified from both developmental processes (Figure 1A, Bai et al., 2017). The ribosomal RNA and tRNA are essential components during mRNA translation. The modulization of both translational components may indicate a coordinated regulation for shaping the translational program during both seed maturation and germination. Redox balancing is another pivotal process for both seed maturation and germination as the desiccation stress imposed low-oxygen stress could lead to an accumulation of electrons from inner mitochondrial membrane, resulting in reactive oxygen species (ROS) accumulation (Patela et al., 2019). ROS in seed is important signaling molecular for breading the seed dormancy while excess of ROS accumulation could trigger ageing process leading to seed deterioration and death (El-Maarouf-Bouteau and Bailly, 2008; Bailly and Kranner, 2011; Leymarie et al., 2012). NAD1A/B, 2A, 3, 4, 5A, 6 dehydrogenases, NADH oxidoreductases and cytochrome b/b6 proteins are specifically identified in this module that are involved in cellular respiration and mitochondrial electron transportation, underlining a translational control of redox balancing during seed maturation and germination.

Large amount of transporter genes are specifically enriched in module 2 which are involved in transporting different ions such as calcium and manganese (ECA3, AT1G10130), nitrate (NRT1.1, AT1G12110) sodium, potassium and chloride (CCC1, AT1G30450), and other small molecules such as purine (AT1G21070), phytochelatin and arsenite (ABCC1, AT1G30400), and xenobiotic transporter (ABCC5, AT1G04120). These transporters are accumulated in transcripts level during seed maturation and consistently translate during the same developmental phases, likely indicating an extensive metabolic exchange during seed reserve filling, which provide the essential substrate for early seed germination. However, the specific function of these translationally associated transporter genes remains to be revealed.

Module 3 contains large number of proteins well known to be important for seed maturation such as LEA proteins, oleosin proteins and protein involved ABI3 cascades, strongly indicating a translational coordination between the protein translated during seed maturation. Specifically, module 3 contains genes that increased in transcript abundance during maturation (Fig 4E), and therefore likely the seed stored mRNAs. As recently reported seed stored mRNAs are important for seed germination by translation at early stage during seed germination (Bai et al., 2019). This is also reflected by the relatively low but dramatically enhanced PO during maturation and germination respectively.

Sequence motives are also grouped according to the network pattern, which further indicating the biological relevance of the SeedTransNet. We focused on one of the identified motifs specific for the seed module - adenosine enriched motif (A-enriched motif) and confirmed its function in vivo. Possibly this sequence represents a signal localized in the transcript 5’UTR directly upstream of start codon that can mediate the initial interaction between the mRNA and RBP or regulatory RNA species. This interaction is then affecting the translational machinery and thus determines the translational efficiency. A similar mechanism is recently identified by the interaction between a poly(A) binding protein and a similar motif in mediating pathogen resistance in Arabidopsis (Xu et al., 2017). The identification of these sequence motifs - regulatory factor pairs will become the prime activity in research concerning translational regulation and the 1800 RBPs of the Arabidopsis genome might represent the same regulatory potential as the 1200 sequence specific DNA binding proteins, the transcription factors (Dedow and Bailey-Serres, 2019).

We also explore the function for one of the central hubs identified from SeedTransNet named *PM19L1*, which was shown to regulate seed dormancy by Barrero et al (2019) during or preparation of the manuscript. We confirm the data from (Barrero et al 2019) and also show that the gene is important for seed longevity (Figure 6A). The high transcript level specifically in dry seeds and its connection with other seed maturation genes indicating its importance during seed maturation. (Figure 6C, Supplemental Figure 7). The PM19L1 homolog was originally identified in wheat (*Triticum aestivum*) as AWPM-19 (ABA-induced Wheat Plasma Membrane Polypeptide-19), which responses to ABA-induced freezing tolerance (Koike et al., 1997). Recently a rice homolog OsPM1 is identified as associated with ABA induced drought tolerance and seed germination speed (Yao et al., 2018). With SeedTransNet, we could further validate its down-stream target *AT2G05580.* PM19L1 is influencing the ribosome association and likely the translation of *AT2G05580*.

In all, the mRNA translation is extensively modulated using several different molecular mechanisms during seed maturation. This manuscript reveals a small part of the regulatory potential. We expect in the future to present further translationally regulatory mechanism during development such as presented in this manuscript but also in response to various stimuli and stress. Plant plasticity is most likely not solely relying on transcriptional regulation. Translational regulation is expected to play a much greater role and what is currently described in the literature. This regulatory potential will open new routes for crop improvement, so desperately needed in times in which our agricultural system are challenged.

## Materials and methods

### Plant material and growth conditions

Maturing seeds were *Arabidopsis thaliana* accession Columbia-0 grown in three biological replicates with 50 plants each. Flowers were tagged at each time point after flower opening and pollination. Seeds were harvested at each specified developmental stage including 12, 15, 18 and 20 days after flowering. About 200-400 mg of seed were harvested and frozen in liquid nitrogen, freeze-dried and stored at −80 °C until further analyzed.

### Isolation of total RNA and polysomal RNA and polysome analysis

Isolation of total and polysomal RNA was performed according to (Bai et al., 2017). Ribosomes were purified using approximately 400 mg of freeze-dried ground tissue extracted with 8 ml of polysome extraction buffer, PEB (0.25 M sucrose, 400 mM Tris, pH 9.0, 200 mM KCl, 35 mM MgCl_2_, 5 mM EGTA, 50 μg/mL Cycloheximide, 50 μg/mL Chloramphenicol). The extracts (10 ml) were loaded on top of an 8 ml sucrose cushion (1.75 M sucrose in PEB) and centrifuged (18h, 90,000 g) using a Beckman Ti70 rotor (Beckman Coulter, Brea, USA). The resulting pellet was resuspended in 0.5 ml suspension buffer (200 mM Tris, pH 9.0, 200 mM KCl, 0.025 M EGTA, 35 mM MgCl_2_, 5 mM DTT, 50 μg/mL Cycloheximide, 50 μg/mL Chloramphenicol) and loaded on a 4.5 ml 20-60% linear sucrose gradient, centrifuged at 190,000 g for 1.5 h, at 4°C using Beckman SW55 rotor (Beckman Coulter). After ultracentrifugation, the gradients were fractionated into 20 fractions using a Teledyne Isco Density Gradient Fractionation System (Teledyne Isco. Lincoln, USA) with online spectrophotometric detection (254 nm). The fractions corresponding to the polysome region in the ribosome profile were pooled for further analysis. The ribosome abundance is reflected by the area under the curve and was calculated after subtracting the baseline obtained by measuring a blank gradient and normalizing to total area under the curve to account for possible uneven loading of the gradients.

### Data analysis

Affymetrix Arabidopsis Gene 1.1 ST Arrays (Affymetrix, Santa Clara, USA) were hybridized using the GeneChip® 3’ IVT Express kit (cat. # 901229) according to instructions from the manufacturer. Hybridization data were analyzed and gene specific signal intensities were computed using the R statistical programming environment (www.R-project.com), the BioConductor package affy (Gautier et al., 2004) and the Brainarray cdf file ver. 17.1.0 (http://brainarray.mbni.med.umich.edu/). DNA microarray data are available in the Gene Expression Omnibus (GEO) repository (http://www.ncbi.nlm.nih.gov/geo/) under accession number GSE127509. The limma and affy package were used for RMA normalization (Irizarry et al., 2003). Probe sets with intensity signals that never exceeded the noise threshold (logExprs<4 in all samples) were removed. A linear Bayesian model were applied for assessing differential expression using the limma package (Smyth, 2004). Principle component analysis (PCA) was performed using TM4 (Saeed et al., 2003).

### GO enrichment analysis

Gene Set Enrichment Analyses were performed using an in-house utility: gopher; an algorithm developed at the Umeå Plant Science Center. For Gene Ontology enrichment, it uses the Parent-Child test (Grossmann et al., 2007) and a Benjamini-Hochberg multiple testing correction. Significant terms were selected if their FDR was <= 0.05.

### Sequence feature analysis

Genes with significantly increased and decreased polysomal occupancy (PO) at each translational shift were compared to all mRNAs expressed in the experiment as background for enrichment of sequence features using custom scripts. Different parts of the CDNAs (CDS, 5’UTR, 3’UTR, and full transcript) were compares specifically. The distributions of sequences length and GC content were evaluated separately for CDS, 5’UTR, 3’UTR, and full transcript. CDSs were also analyzed for GC3 content, measured using the Effective Number of Codons (Nc) index (Sun et al., 2013), after removing sequences lacking start codons and/or containing premature stop codons and CDSs shorter than 100 codons. The same analyses were performed separately for the CDS of protein-coding genes having both annotated UTR and lacking proper annotation (UTRs called present when length exceeding 1 nucleotide). Given the non-normality of the values distributions, a Wilcoxon signed-rank test was adopted for all statistical comparisons (median as test statistic).

### RNA structural analysis

Experimentally determined structure scores per nucleotide, as provided by (Li et al., 2012), were used to calculate average structure scores of the genes with significantly increased and decreased ribosomal association at each developmental switch. Relative scaling was achieved by averaging the structure scores per region (5’UTR, CDS and 3’UTR) in 100 bins. Standard errors and Student’s t-tests were performed using the Python SciPy module (http://www.scipy.org/).

### Motif analysis

DNA motif analyses were performed using the MEME suite (Bailey et al., 2009), for full transcript, 5’UTR, CDS and 3’UTR sequences, extracted from the TAIR10 database (http://www.arabidopsis.org/). The minimum and maximum motif width was set to 6 and 10, respectively. If a gene had multiple transcripts, only the TAIR10 representative splice form was used. Background dinucleotide frequencies were provided separately for each sequence type. To test specificity of the resulting motifs, FIMO (Bailey et al., 2009) was used to scan all genes represented on the microarray for motif hits in the corresponding sequence type. Motifs with *P*-value lower than 0.001 were considered as significant hits. Obtained motif counts were used to compute the enrichment *P*-value for the gene lists versus the all mRNAs expressed in the experiment as background by means of a one-tailed Fisher’s exact test, performed with a custom script and the R software package (http://www.r-project.org/). The occurrences for Table 1 were derived from the same FIMO outputs. For each motif, the positions on the transcripts, as provided by the FIMO output, were used to calculate the relative number of motifs per (relative) position along the mRNA. Relative scaling was performed in a similar fashion as for the structure scores.

### Data collection and network construction, inference and visualization

In order to collect dataset for a full translational dynamics during seed maturation and germination, DNA microarray data from (Bai et al., 2017) are retrieved in the Gene Expression Omnibus (GEO) repository (http://www.ncbi.nlm.nih.gov/geo/) under accession number GSE65780 (http://www.ncbi.nlm.nih.gov/geo/query/acc.cgi?token=urelcaiqjnqbrux&acc=GSE65780) and GSE127509 (https://www.ncbi.nlm.nih.gov/geo/query/acc.cgi?acc=GSE127509) from current study. The limma and affy package were used for RMA normalization (Irizarry et al., 2003) with R version 3.5.2. Only the genes that passed the noise filter (Log2 > 4) in at least one developmental stage on either transcriptional or translational level were selected for further data processing. Polysome occupancy was calculated by the log2 ratio of polysomal RNA and total RNA according to Bai *et al*., (2017). Polysome occupancy profile with three biological replicates were clustered using Euclidean distance and the visualized with heat map in R by package “pheatmap” (Kolde, 2018). The number of clusters were determined by R package “NbCluster” with “fviz_nbclust” function (Charrad et al., 2015).

Replicated polysome occupancy dataset was used as input for network construction. For network inference, the *seidr* toolkit Seidr was used for meta-network construction (Schiffthaler et al., 2018). In brief, nine gene network inference methods were run: ANOVA (Kuffner et al., 2012), Aracne2, CLR (Faith et al., 2007a), GeneNet (Opgen-Rhein and Strimmer, 2007), GENIE3 (Huynh-Thu et al., 2010), NARROMI (Zhang et al., 2013), Pearson, Spearman, and a modified implementation of TIGRESS (Haury et al., 2012); and their results aggregated into a consensus network using the Top1 method (Hase et al., 2013). An assessment of the scale-free property of the consensus network, fitting a heavy-tailed distribution using a log-log linear model was performed at several candidate thresholds in the range 0.9999 to 1 with a step size of 1e-5. The network transitivity (or average cluster criterion ACC) was calculated within the same range. The best threshold was chosen at 0.99997 with an R^2 scale free fit of 0.95 and an ACC of 0.4. The filtered network were visualized and processed using the Gephi (https://gephi.org/). The network was further partitioned using Infomap (Rosvall and Bergstrom, 2008).

### Seed phenotyping

Seeds were harvested in bulk from four plants for each biological replicate, four biological replicates for each genotype. Germination experiments were performed as described previously (Joosen et al., 2010). In brief, two layers of blue germination paper were equilibrated with 50 ml demineralized water in plastic trays (15×21cm). Six samples of approximately 50 to 150 seeds were spread on wetted papers using a plastic mask to ensure accurate spacing. Piled up trays were wrapped in a closed transparent plastic bag. The experiment was carried out in a 22°C incubator under continuous light (143 μmol m^-2^ s^-1^). Pictures were taken twice a day for a period of 6 d using the same camera and Germinator package software according to (Joosen et al., 2010) for scoring number of seeds that are germinating. Germination was scored using the. To quantify seed dormancy (DSDS50: days of seed dry storage required to reach 50% germination), germination tests were performed weekly until all seed batches had germinated to at least 90% of the seeds. A generalized linear model with a logit link as described (Hurtado et al., 2012) was adapted to calculate DSDS50. Germination data were adjusted by choosing n = 100 and fitted as one smooth curve per line. The observed germination proportion was re-interpreted as having observed y ‘successes’ in n binomial trials (e.g. 75% germinated means y = 75 out of 100 possible ‘trials’). DSDS50 is the closest time point to where a horizontal line at y = 50 crosses the fitted curve. To measure seed longevity, an artificial ageing test was performed by incubating seeds above a saturated ZnSO4 solution (40°C, 85% relative humidity) in a closed tank with circulation for 5-9 d (ISTA, 2012). The seeds were then taken out and germinated on demineralized water as described previously.

### Motif reporter constructs and infiltration

The motif reporter construction was based on the GreenGate system (Lampropoulos et al., 2013). In general, different GreenGate entry vectors were used including pGGA004 with 35S promoter, pGGB003 with a Dumy sequence, pGGC000 empty vector with either a 10 bp poly(A) motif identified from the network module or a 10 bp control motif designed from (http://www.faculty.ucr.edu/~mmaduro/random.htm) and synthesized (Eurofin Scientific, Brussels) respectively, pGGD000 empty vector with luciferase reporter gene cloned from PUGW35 vector, pGGE001 with the RuBisCO gene terminator, and pGGF007 with the Kanamycin resistance gene cassette. All entry vectors were incorporated into the destination vector pGGZ001 according to (Lampropoulos et al., 2013). The final destination vector was sequenced and transformed into the *Agrobacterium tumefaciens* strain (GV3010 pMP90 pSoup) and infiltrated into *Nicotiana benthamiana* leaves. The infiltrated leaves were trimmed after different days of infiltration, sprayed with D-luciferin (Duchefa) solution (1ml firefly D-luciferin, sodium-salt, Duchefa, 0.01% Tween 80). Imaged of luciferase activity is recorded with an exposure time of 10-15 min using a Pixis 1024 (1024×1024) camera system (Princeton Instruments) equipped with a 35 mm, 1:1.4 Nikon SLR camera lens fitted with a DT Green filter ring (Image Optics Components Ltd.) to block delayed auto-fluorescence chlorophyll emissions. Image of the luciferase activity are presented as false color scores (blue indicating low activity and red indicating high activity). After imaging, the infiltrated area of the leaves was excised and snap-frozen with liquid nitrogen and stored in −80° C for further RNA isolation. mRNAs are extracted with TriPure Isolation Reagent (Roche, Basel, Switzerland) according to the total mRNA isolation protocol from (Bai et al., 2017).

## Acknowledgements

We thank Service XS, Leiden, The Netherlands for performing the microarray hybridizations. This work was supported by the Netherlands Organization for Scientific Research. We thank Bio4Energy, a Strategic Research Environment appointed by the Swedish government, for supporting this work. We thank Bastian Schiffthaler from Umeå Plant Science Center for the support with the network analysis.

## Author contributions

B.B., J.H. and L.B. designed research; B.B. performed research; B.B., N.D. and S.H. analyzed data; B.B., J.H. and L.B. wrote the paper.

## Supplemental Data

Supplemental Figure 1. Transcriptional profile during seed maturation.

Supplemental Figure 2. Comparison of translationally regulated genes during seed maturation with other set of translationally regulated genes.

Supplemental Figure 3. Intensity profiles of genes with changed PO.

Supplemental Figure 4. Sequence features of genes translational regulated during seed maturation.

Supplemental Figure 5. Threshold selection for SeedTransNet construction.

Supplemental Figure 6. Comparison between regulons identified in SeedTransNet and previous published regulons during seed maturation.

Supplemental Figure 7. Network database comparison for genes predicted from SeedTransNet.

Supplemental Figure 8. Complementation of *atpm1* seed longevity phenotype by overexpression of the AtPM1 gene.

Supplemental Table 1. Raw normalized intensity levels of total RNA (T) and polysomal RNA (P) (log2 transformed) during seed maturation.

Supplemental Table 2. Translational regulated genes during seed maturation

Supplemental Table 3. GO enrichment analysis for the translationally regulated genes during seed maturation

Supplemental Table 4. GO term enriched in the top 100 highest CDS length, structure score in the transcript increased in PO during seed maturation and the associated genes

Supplemental Table 5. Average Polysomal occupancy (PO) of transcripts in the hierarchical clusters of the combined time course data of polysome occupancy during seed maturation and germination

Supplemental Table 6. GO enrichment analysis of the individual clusters from hierarchical clustering of PO time course during seed maturation and germination.

Supplemental Table 7. Seed translation net (SeedTransNet). Columns listed are resource, target, linkage weight, direction for the network.

Supplemental Table 8. Infomap community prediction and gene linkage inference of SeedTransNet.

Supplemental Table 9. GO enrichment analysis of genes in SeedTransNet modules

Supplemental Table 10. Genes identified in from chromosome 2 and mitochondria in M1 and their enriched functions.

Supplemental Table 11 Genes included in the LEA protein network as identified in the Module 3 of SeedTransNet.

Supplemental Table 12. Genes included in the ABI3 core regulon network as identified in Module 3 module of SeedTransNet.

Supplemental Table 13. Raw normalized signals in maturation dataset of the known seed maturation related genes. Column represents the gene ID, gene name, direction of translational regulation, type of translational subgroup (with gene number), and the expression pattern for both polysomal RNA (P) and total RNA (T) that isolated from 12, 15, 18 and 20 days after flowering (DAF) respectively.

