## Supplemental Figure for "Transcriptome and translatome profiling and translational network analysis during seed maturation reveals conserved transcriptional and distinct translational regulatory patterns"

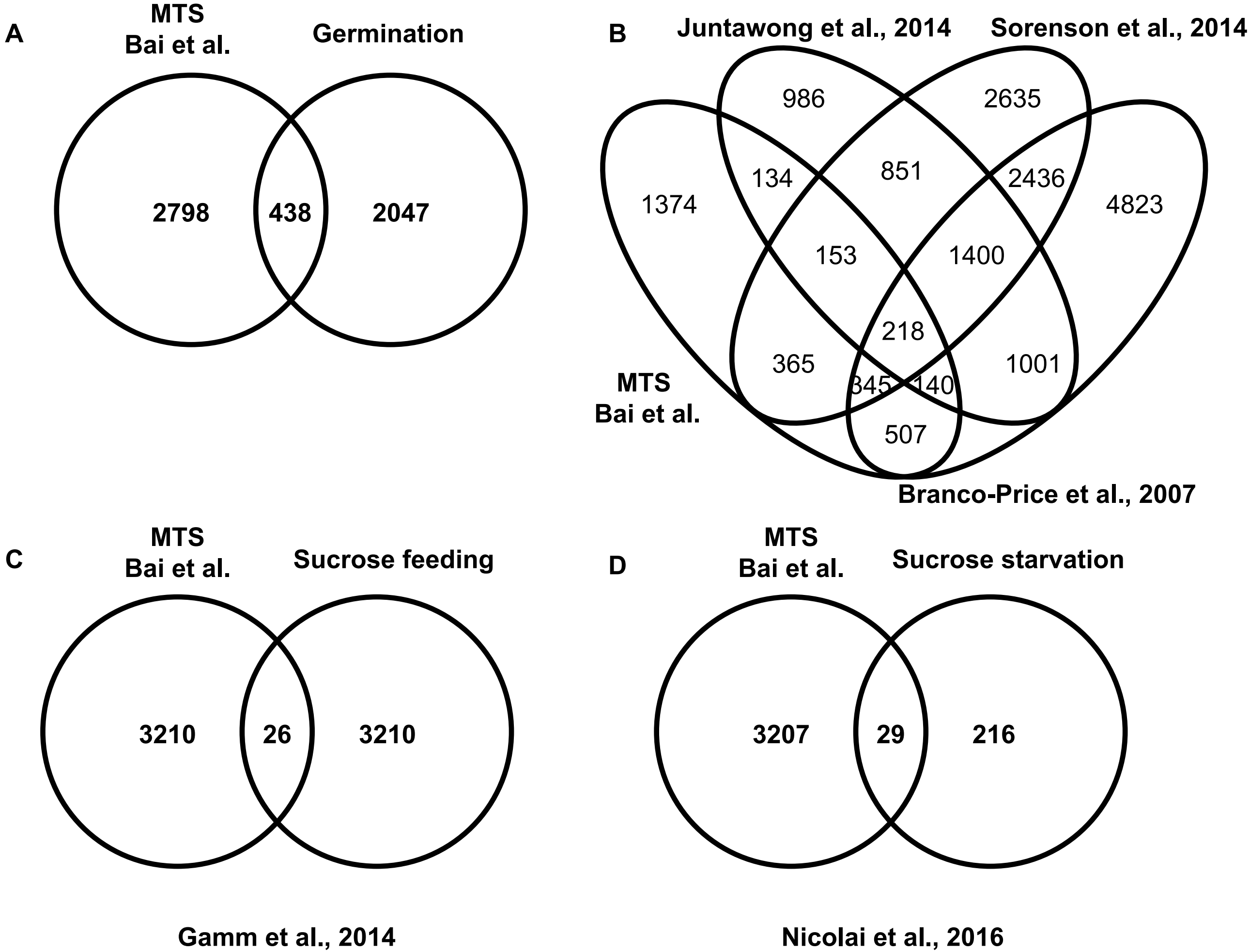


Supplemental Figure 1

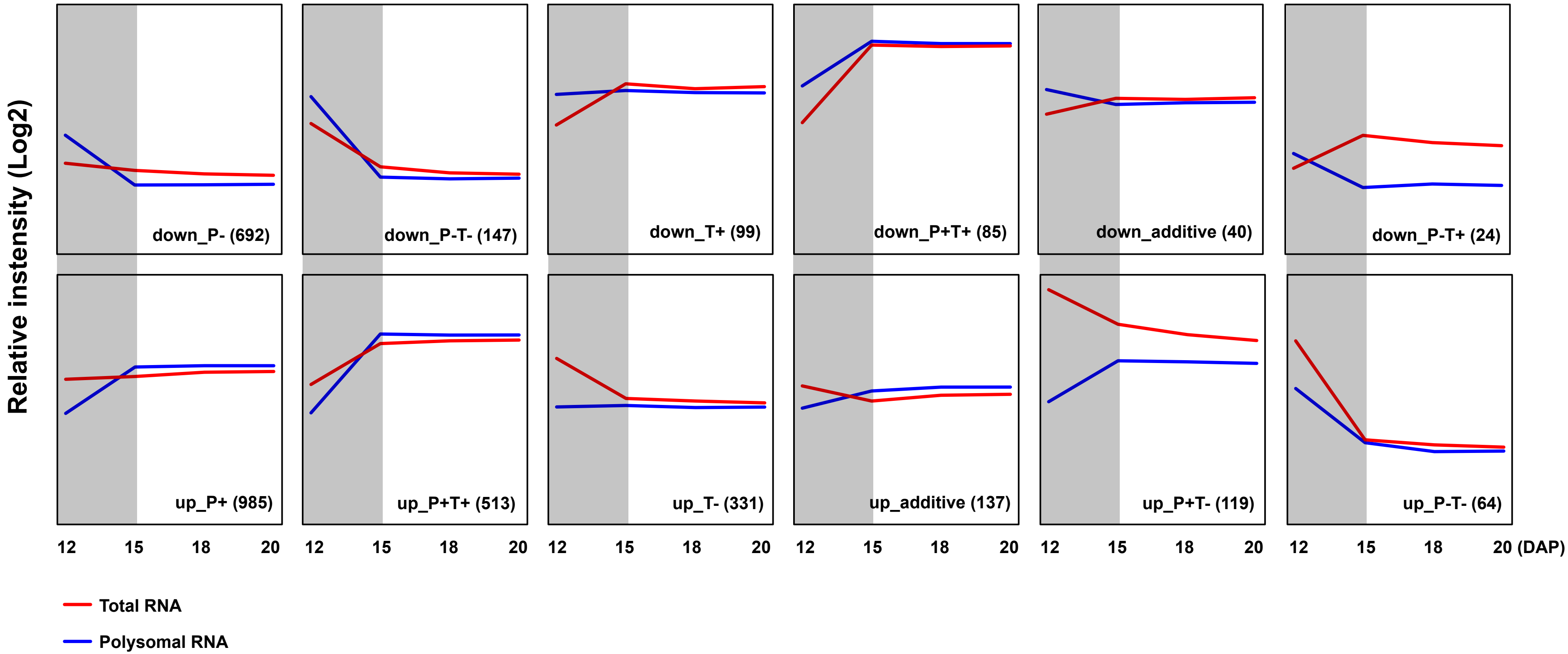


**Supplemental Figure 2**

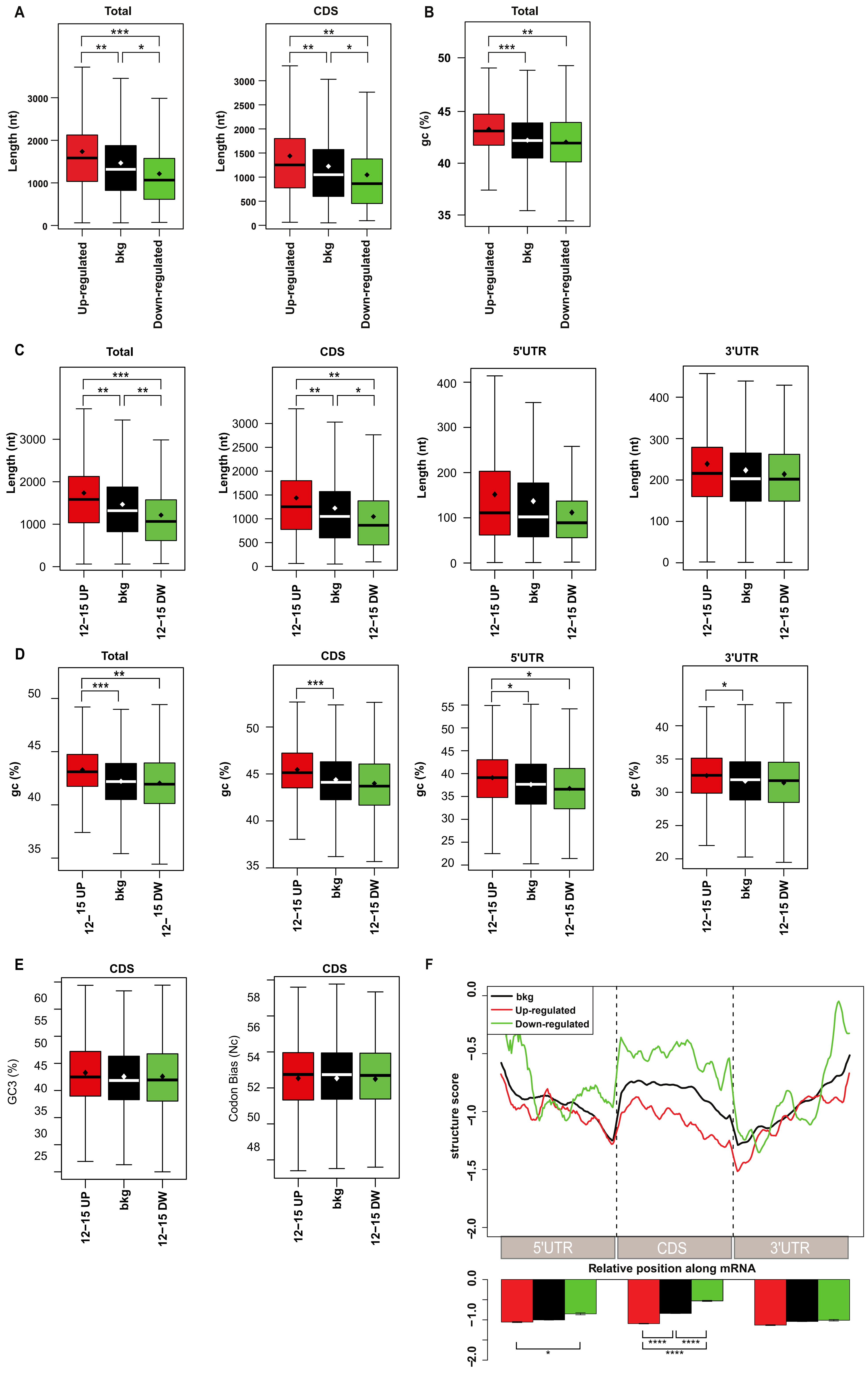


**Supplemental Figure 3**

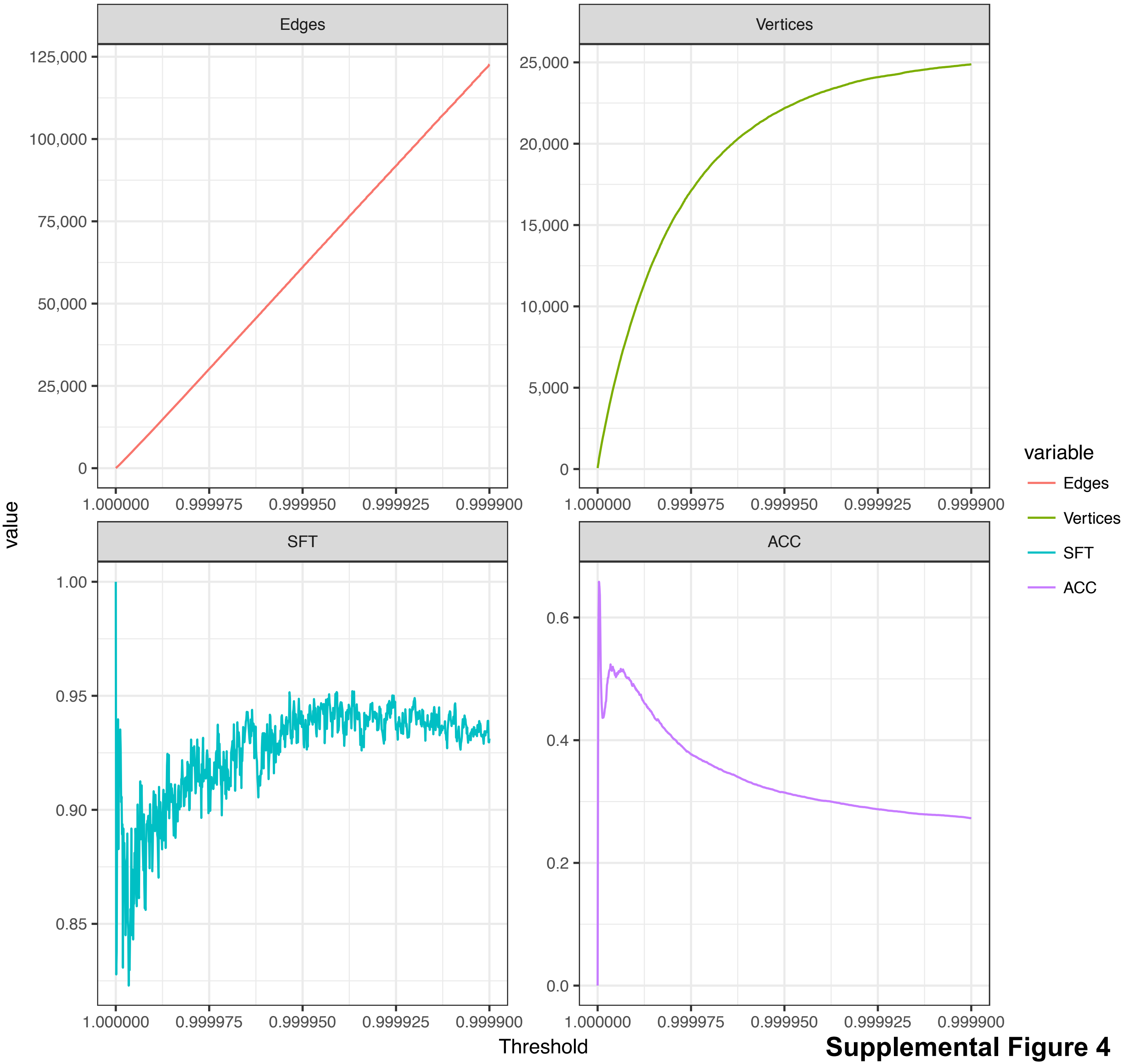


**Supplemental Figure 4**

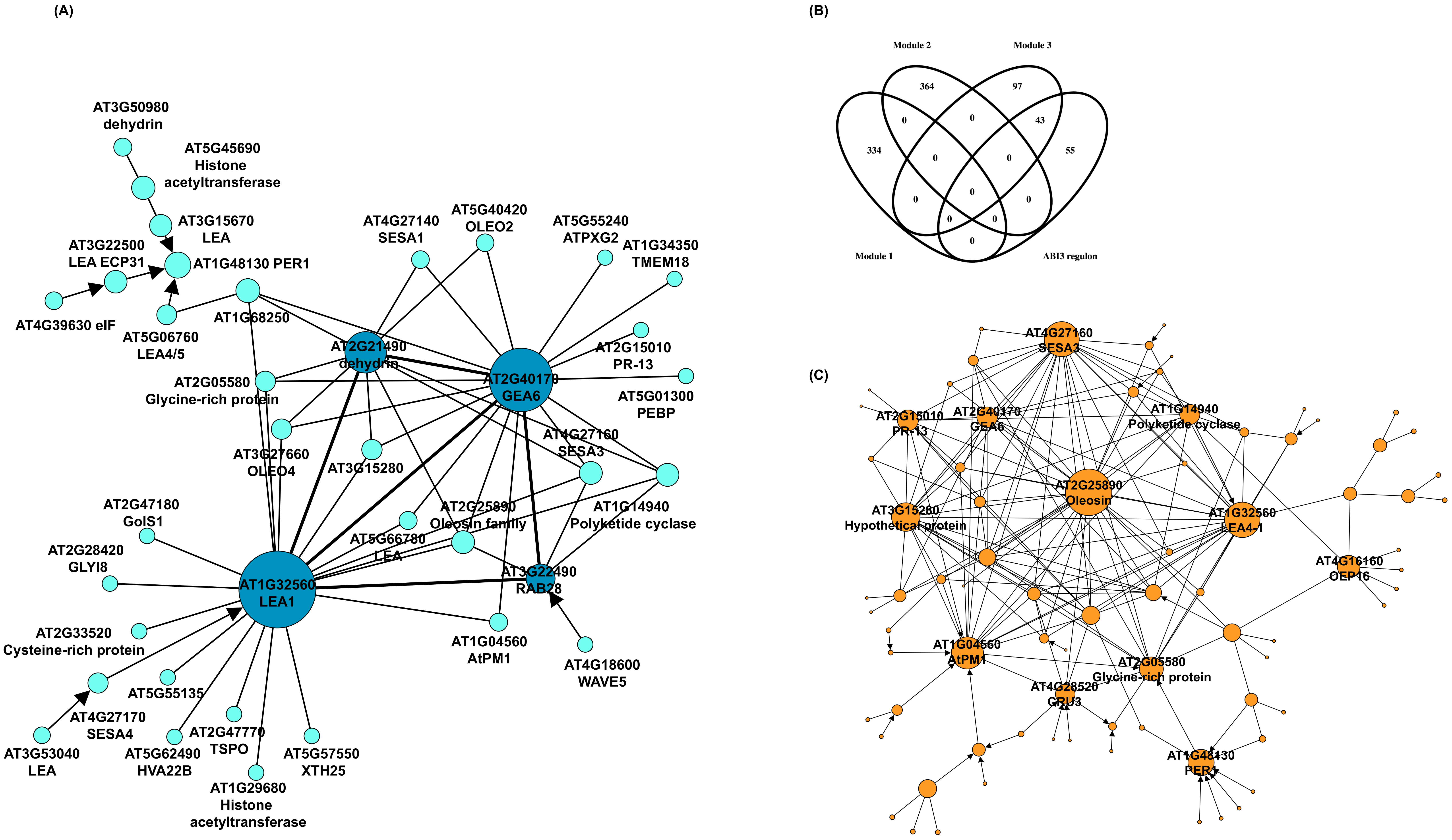


Supplemental Figure 5

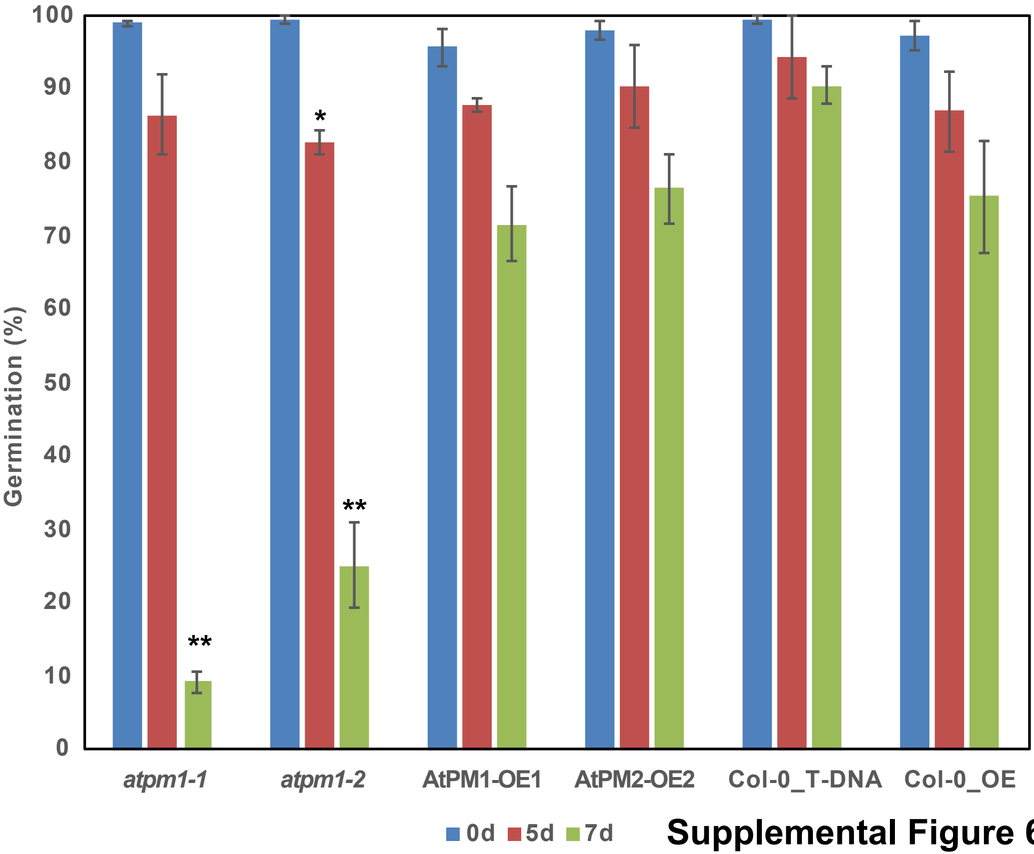


**Supplemental Figure 6**

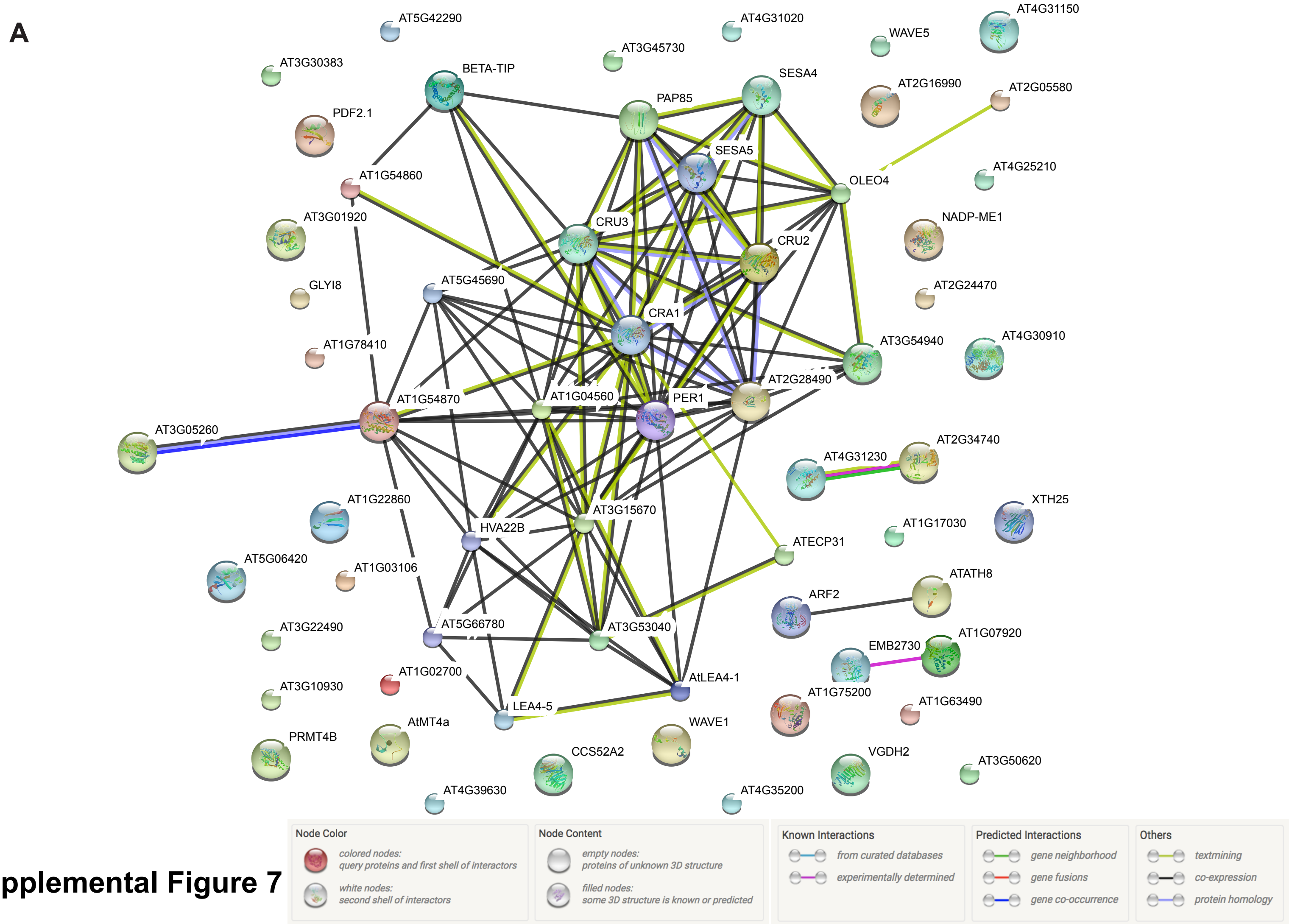


**Supplemental Figure 7**

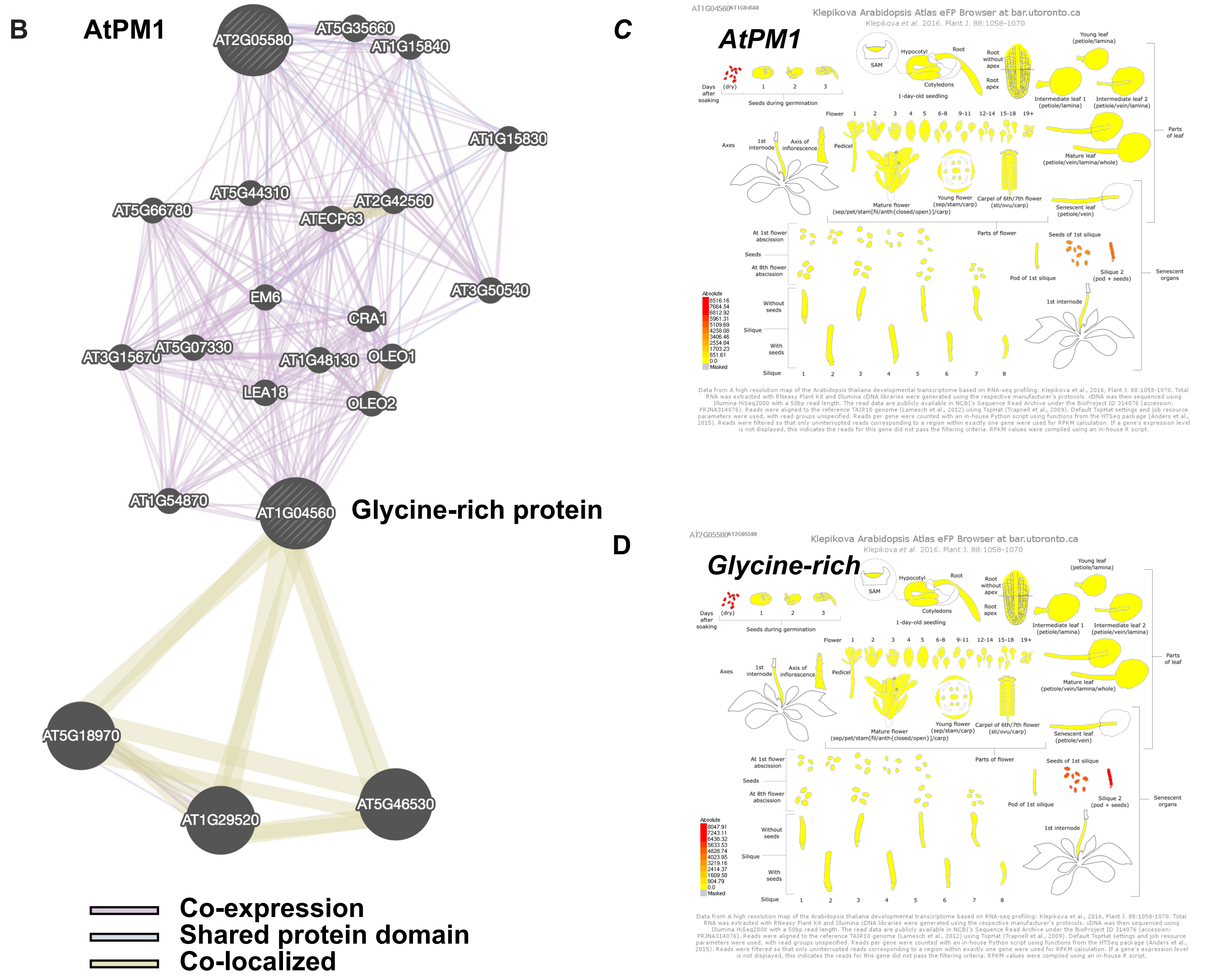


**Supplemental Figure 8**

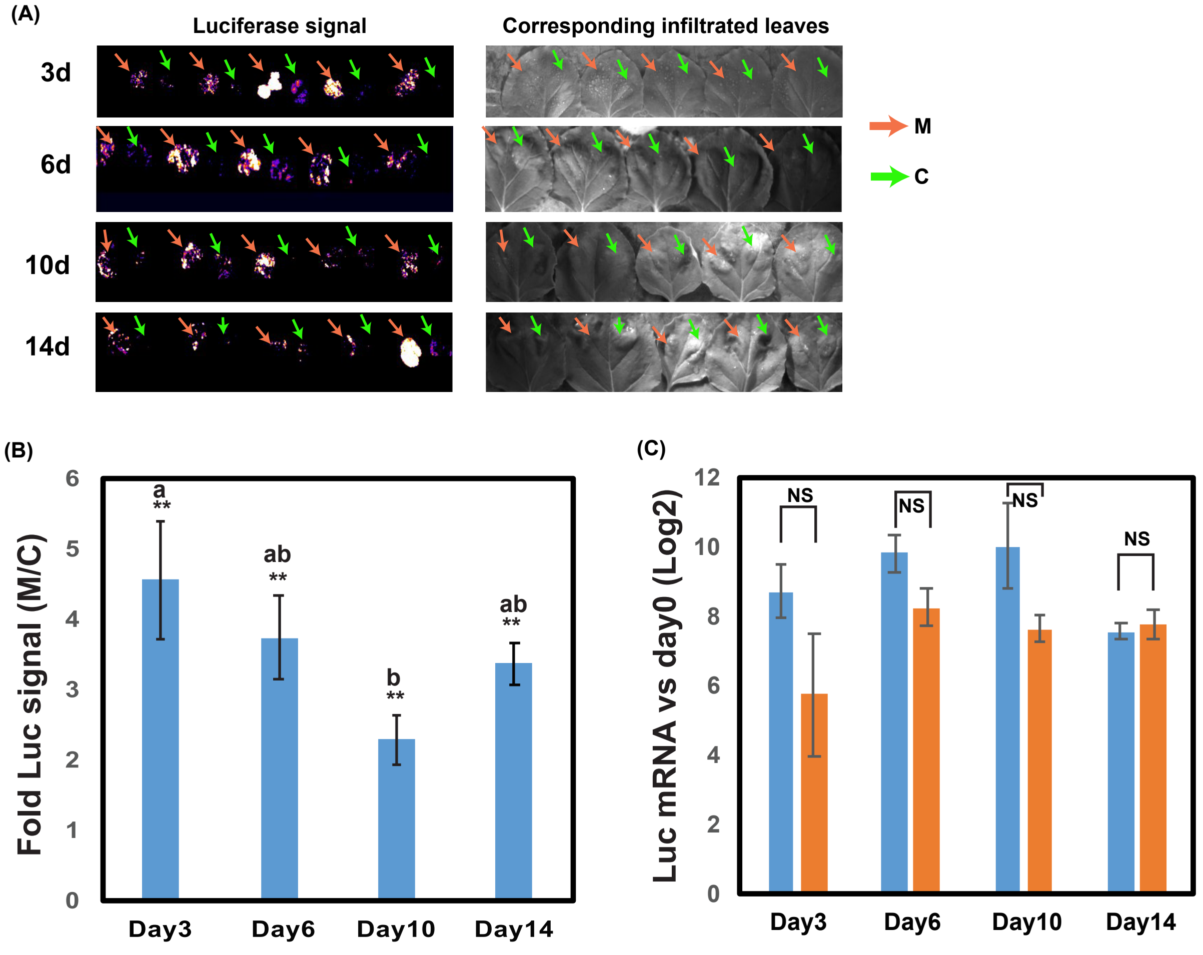


**Supplemental Figure 9**
